# Hybridization and its impact on the ontogenetic allometry of skulls in macaques

**DOI:** 10.1101/2023.04.19.536293

**Authors:** Tsuyoshi Ito, Ryosuke Kimura, Hikaru Wakamori, Mikiko Tanaka, Ayumi Tezuka, Atsushi J. Nagano, Yuzuru Hamada, Yoshi Kawamoto

## Abstract

The role of hybridization in morphological diversification is a fundamental topic in evolutionary biology. However, despite the accumulated knowledge on adult hybrid variation, how hybridization affects ontogenetic allometry is less well understood. Here, we investigated the effects of hybridization on postnatal ontogenetic allometry in the skulls of a putative hybrid population of introduced Taiwanese macaques (*Macaca cyclopis*) and native Japanese macaques (*M. fuscata*). Genomic analyses indicated that the population consisted of individuals with varying degrees of admixture, formed by male migration from Japanese to Taiwanese macaques. For overall skull shape, ontogenetic trajectories were shifted by hybridization in a nearly additive manner, with moderate transgressive variation observed throughout development. In contrast, for the maxillary sinus (hollow space in the face), hybrids grew as fast as Taiwanese macaques, diverging from Japanese macaques, which showed slow growth. Consequently, adult hybrids showed a mosaic pattern, i.e., the maxillary sinus is as large as that of Taiwanese macaques, while the overall skull shape is intermediate. Our findings suggest that the transgressive variation can be caused by prenatal shape modification and non-additive inheritance on regional growth rates, highlighting the complex genetic and ontogenetic bases underlying hybridization-induced morphological diversification.

## 1. Introduction

A growing understanding indicates that interspecific hybridization is not rare, even in animals (Mallet 2005; Taylor and Larson 2019). Furthermore, hybridization is not always maladaptive and can facilitate evolutionary novelty and diversity by generating gene combinations not achieved in the parental lineages (Seehausen 2004; Arnold and Kunte 2017; Kagawa and Takimoto 2018). Thus, hybridization must have played a critical role in the evolution of various taxa and has consequently been of significant interest to evolutionary biologists.

Primate hybridization is of particular interest because it enhances our understanding of the effects of archaic human gene introgression on the modern human lineage. Molecular studies show that hybridization is pervasive in primates, with relics of ancient hybridization or the existence of a natural hybrid zone identified in various taxa of primates, e.g., macaques, baboons, vervet monkeys, marmosets, colobine monkeys, and howler monkeys (Roos et al. 2011; Malukiewicz et al. 2014; Cortés-Ortiz et al. 2015; Svardal et al. 2017; Rogers et al. 2019; Osada et al. 2021). Furthermore, several studies have assessed the morphological effects of hybridization in nonhuman primates (Ackermann et al. 2019). In baboons and gorillas, hybrids exhibit anomalies such as supplementary rotated teeth or atypical cranial sutures at higher frequencies than in their parental lineages (Ackermann and Bishop 2010; Ackermann et al. 2014). In various taxa of primates, hybrids often show heterotic/transgressive phenotypes or increased variation in cranial and postcranial morphology (Cheverud et al. 1993; Ackermann et al. 2014; Fuzessy et al. 2014; Ito et al. 2015; Eichel and Ackermann 2016; Boel et al. 2019; but see Buck et al. 2021). Such hybridization-driven genetic perturbations may contribute to the morphological diversity in primate evolution.

Any morphological evolution is achieved by the modification of the ancient ontogenetic trajectory. For example, the distinct adult human craniofacial morphology is formed by divergence from the hominid common ontogenetic trajectory in the early postnatal period (Mitteroecker et al. 2004). The difference in brain (endocast) shape between modern humans and Neanderthals was also formed by early divergence (Gunz et al. 2010); however, the overall craniofacial difference between the two species was formed by a more or less parallel (but not identical) ontogenetic trajectory that originated from the established difference at birth (Ponce de León and Zollikofer 2001; Bastir et al. 2007). Because the ontogenetic trajectory reflects the expression of genes that control morphology during development, hybridization-induced genomic admixture potentially alters an ontogenetic trajectory. Hybrid Sulawesi macaques are reported to show acceleration (slope increase) and postdisplacement (intercept decrease) in the ontogenetic trajectory of several craniofacial traits, although this dissociation does not account for adult heterosis (Schillaci et al. 2005). Predisplacement (intercept increase) occurs in the ontogenetic trajectory of body weight in inter-subspecies male hybrids of rhesus macaques but is lacking in female hybrids (Smith and Scott 1989; Taylor and Schillaci 2008). In non-primate taxa, several studies have reported different types of ontogenetic modifications, such as pure heterochrony, directional changes, or intermediate but maternally biased trajectories, in the head or body shape of fish hybrids (Holtmeier 2001; Corse et al. 2012; Sinama et al. 2013; Santos-Santos et al. 2021). However, with the exception of these studies, knowledge of the effects of interspecific hybridization on ontogenetic trajectories remains limited, especially in mammals.

Anthropogenic hybridization provides an opportunity to study the consequences of hybridization, although it is controversial in the context of conservation. In the Oike area of Wakayama Prefecture, Japan, introduced *Macaca cyclopis* (Taiwanese macaques) that escaped from a zoo in the 1950s hybridized with native *M. fuscata* (Japanese macaques) (Ohsawa et al. 2005). By 2004, the population had grown to approximately 300 individuals with four troops (Ohsawa et al. 2005). Genetic analyses suggested that the Oike hybrid population was formed by male immigration into *M. cyclopis* from the neighboring native population of *M. fuscata* (Kawamoto et al. 2001, 2008a). To prevent genetic introgression and its impacts on the ecosystems, the government began the elimination of introduced and hybrid individuals from the Oike area in 2002 and finally declared eradication in 2017, with a total of 366 individuals eradicated from the Oike area (Shirai et al. 2018). The Oike hybrids show several intriguing morphological variations. For example, unlike previous studies on hybrids of other primate taxa, the Oike hybrids did not show a marked increase in dental anomalies or craniofacial variability compared to their parental species (Boel et al. 2019). Relative tail length variation followed the expectations of the additive model (Hamada et al. 2012). Adult craniofacial shape followed additivity or was slightly closer than expected to that of *M. fuscata*; in contrast, maxillary sinus size (hollow space in the face) was much closer to that of *M. cyclopis* than additivity (Ito et al. 2015; Boel et al. 2019). However, it remains unclear how these adult differences manifested during development.

This study aimed to elucidate the effects of hybridization on the ontogenetic trajectory of craniofacial morphology using samples from the Oike hybrid population. First, we estimated the population structure of the Oike population as well as the hybrid index and class (generation) of individuals based on genome-wide single nucleotide polymorphisms (SNPs). Second, we compared morphological variation and ontogenetic trajectories of the cranium, mandible, and maxillary sinus with the hybrid index and among hybrid classes. Comparing ontogenetic trajectories among hybrids and parental lines will improve our understanding of the genetic and ontogenetic basis of hybridization-induced morphological diversification.

## 2. Materials and methods

### 2.1. Samples

Almost all available samples from the Oike hybrid population were screened, including 245 skeletal specimens and 287 DNA samples (Table S1; Fig. 1). Note that some samples were filtered out prior to the main analyses (see below for details). Of these, the 215 skeletal specimens could be linked to DNA samples, where we linked them as far as the IDs on the records could be uniquely identified, even if they did not show an exact match. These samples came from collections obtained under the government eradication program between 2003 and 2006; no animals were sacrificed for research purposes. Skeletal specimens are stored at the Primate Research Institute (now the Center for the Evolutionary Origins of Human Behavior), Kyoto University (Inuyama, Japan). Note that individuals judged to be pure Japanese macaques in the Oike area had been released (Hamada et al. 2008).

**Figure 1:**
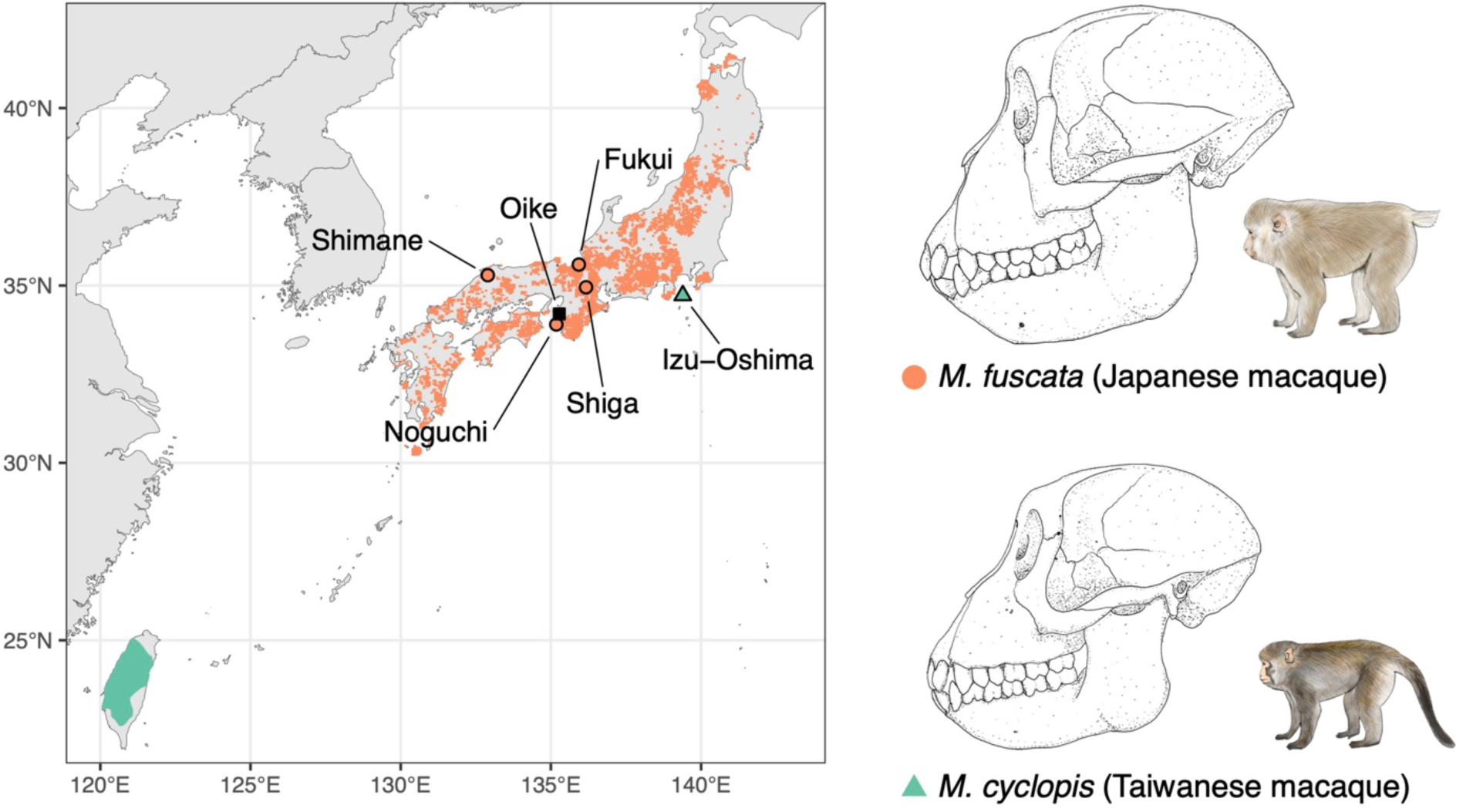
Natural distribution of *Macaca cyclopis* (Taiwanese macaques; blue) and *M. fuscata* (Japanese macaques; magenta) and sampling sites. Oike is the location of the hybrid population. *M. fuscata* skeletal specimens are from Fukui (*N* = 15), Shimane (*N* = 21), Shiga (*N* = 49), and Noguchi (*N =* 1). *M. cyclopis* skeletal specimens are from Izu-Oshima Island (*N* = 20), where introduced *M. cyclopis* is distributed. This map was created by the authors using IUCN Red List Data (version 3, May 2017) and distribution surveys of Japanese animals (mammals) by the Biodiversity Center of Japan, Nature Conservation Bureau, Ministry of the Environment.

In addition, to test whether pure parental individuals remained in the Oike population, we examined DNA from pure captive *M. cyclopis* (*N* = 4) and *M. fuscata* (*N* = 4) housed at the Primate Research Institute. The *M. fuscata* individuals were descendants of individuals transferred from Arashiyama, Kyoto, approximately 100 km north-northeast of the Oike area. There is no record of the exact origin of the *M. cyclopis* individuals. Sample collection from these captive individuals was performed according to the recommendations of the Guidelines for Care and Use of Nonhuman Primates Version 2 and 3 of the Primate Research Institute, Kyoto University (2002, 2010). DNA was extracted from blood using the phenol-chloroform method. Skeletal specimens of *M. cyclopis* (*N* = 54) and *M. fuscata* (*N* = 88) stored at the Primate Research Institute and the Japan Monkey Centre (Inuyama, Japan) were used as surrogates for parental individuals for morphological analyses (because the actual parental populations are unknown). Captive specimens and specimens of unknown origin were initially included, but preliminary analyses revealed morphological differences between wild and captive specimens (see below). Therefore, only wild specimens (*N =* 18 for *M. cyclopis*, *N =* 86 for *M. fuscata*) were used for the main analyses. Wild *M. cyclopis* specimens were obtained from the introduced population on Izu-Oshima Island, Tokyo, Japan, where *M. fuscata* is not distributed. Wild *M. fuscata* specimens were obtained from localities around the Oike area and are therefore likely to be genetically close to a parental population (Suzuki-Hashido et al. 2015; Ito et al. 2021).

### 2.2. Genetic analyses

Double-digest restriction site-associated DNA sequencing was performed based on Peterson’s protocol (Peterson et al. 2012) with modifications (Sakaguchi et al. 2015, 2017). After filtering raw reads for overall sequence quality, genotypes were called using STACKS 2.59 (Rochette and Catchen 2017). Because STACKS provides diploid genotypes, genotypes on male sex chromosomes were converted to haploid using diploid2haploid.py (https://github.com/itots/diploid2haploid.git), omitting heterogeneous genotypes (1.2% for the X chromosome and 0.8% for the Y chromosome). After filtering, we obtained 6461 variant sites in 289 individuals for the autosome (total genotyping rate = 0.96), 171 sites in 289 individuals for the X chromosome (0.96), and 12 sites in 152 individuals for the Y chromosome (0.95). See Supplementary Note for details.

Global ancestry was inferred from the SNPs separately for autosomes and X chromosomes using ADMIXTURE (Alexander and Novembre 2009); the male X chromosome was treated as haploid using the --haploid flag. Principal component analysis (PCA) was performed using the “adegenet” package (Jombart 2008) in R (R Developmental Core Team 2021). For the Y chromosome, *K*-means cluster analysis (*K* = 2) was performed based on the first principal component (PC). Six samples showed inconsistencies in global ancestry or sex identification with data from Kawamoto et al. (2008a) and were removed from the subsequent analyses. See Supplementary Note for details.

Individuals in the Oike population were classified into 23 *M. cyclopis* (*Q* < 0.001), 11 *M. fuscata* (*Q* > 0.999), and 242 putatively admixed individuals (0.001 ≤ *Q* ≤ 0.999) based on the ADMIXTURE *Q* score (*K* = 2) of autosomal SNPs. Diagnostic markers were detected as differentially fixed sites (allele frequency difference between the two parental populations [δ] = 1), omitting loci with missing genotype rates within each of the two parental populations > 0.1. Hybrid index, the proportion of ancestry belonging to *M. fuscata*, and interspecific heterozygosity, the proportion of loci with alleles from both parental populations, were calculated from the 196 autosomal diagnostic markers using a custom R script.

Hybrid classes were estimated for individuals in the Oike population using NewHybrids 1.1 (Anderson and Thompson 2002). This Bayesian-based method calculates the posterior probability that an individual belongs to a genotype frequency class. The SNP dataset of 196 autosomal diagnostic markers was converted into a NewHybrid format using PGDSpider 2.1.1.5 (Lischer and Excoffier 2012). The “z” option was used to instruct NewHybrid that individuals with a hybrid index = 0 or 1 were pure parental individuals. Ten replicates were run with randomly differentiated seeds using the “parallelnewhybrid” R package (Wringe et al. 2017), with minor modifications (https://github.com/itots/parallelnewhybrid.git). Each run started with Jeffery’s prior and six default genotype frequency classes and consisted of 10,000 Markov chain Monte Carlo (MCMC) sweeps after 10,000 burn-in iterations. We confirmed MCMC convergence by visual inspection. Based on the mean posterior probabilities, each individual was assigned to one of six genealogical classes: *M. cyclopis* (P0), *M. fuscata* (P1), F1, F2 hybrids, and backcrosses of *M. cyclopis* (BC0) and *M. fuscata* (BC1). We discarded assignments with mean posterior probability <0.99, P0 with hybrid index >0, P1 with hybrid index <1, and F1 with interspecific heterozygosity <0.95; these were removed from analyses examining hybrid class differences (shown as NA in figures). Note that individuals identified as F2, BC0, and BC1 may include later generations; unfortunately, it is difficult to distinguish between hybrid classes of recombinant generations based on unphased and limited numbers of variants.

### 2.3. Morphological analyses

The 10-step developmental phase was determined based on the dental eruption pattern and spheno-occipital synchondrosis, according to Mouri (1994), with minor modifications (Table S2). Based on this, developmental stages were determined as infant (developmental phase <3), juvenile (<8), and adult (≥8). These stages were used to evaluate age structure and developmental changes in morphology.

The cranium and mandible were scanned by computed tomography (CT), and 3D landmarks were digitized (cranium: number of landmarks = 50, mandible: 28, and maxillary sinus: 10) using 3D Slicer (https://www.slicer.org/) (Kikinis et al. 2014) (Table S3; Fig. 2). See Supplementary Note for details. All landmarks on the maxillary sinus were type III and were used only to evaluate size, not shape. Digitization was performed twice on all specimens by a single observer to evaluate and minimize measurement errors. Specimens with a missing landmark ratio > 0.2 (three specimens for the cranium and one for the mandible) were removed. For the remaining specimens, missing landmarks (1–10 in 53 specimens for the cranium, 1–4 in 10 specimens for the mandible, and 1–2 in 30 specimens for the maxillary sinus) were imputed based on the Bayesian PCA method using the “LOST” R package (Arbour and Brown 2014, 2020).

**Figure 2:**
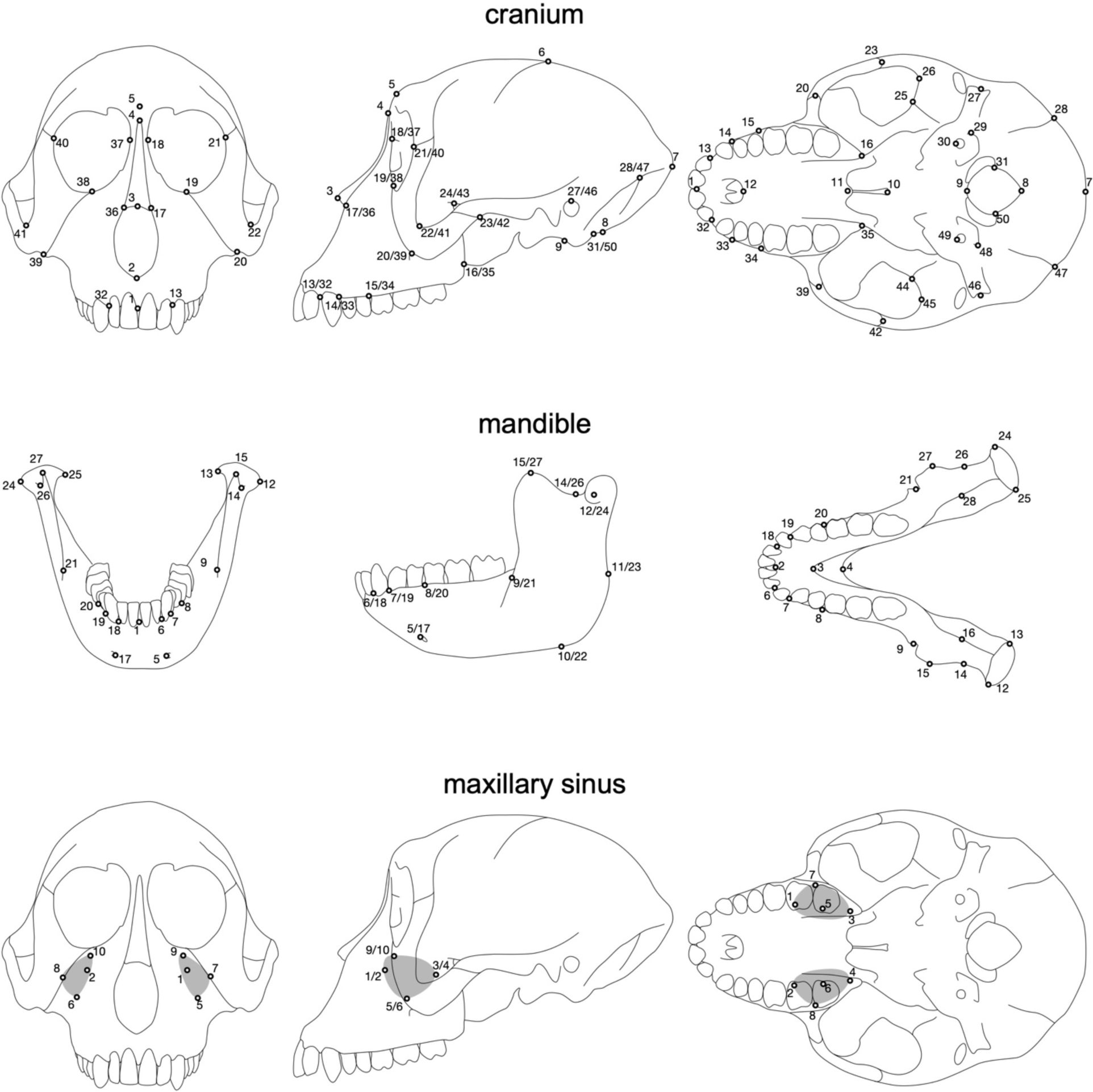
Landmarks used in this study.

Generalized Procrustes analysis and Procrustes analysis of variance were performed to assess measurement error using the “geomorph” R package (Collyer and Adams 2018, 2021; Adams et al. 2021; Baken et al. 2021). Here, centroid size was calculated as the square root of the sum of the squared distances of a set of landmarks from their centroid. For the cranium and mandible, shape measurement errors were negligible compared to inter-individual variation (Table S4), and a symmetric component was used in the following analyses. Measurement errors were negligible for the centroid size of the cranium, mandible, and maxillary sinus (Table S5), and the mean of replicates (mean of replicates and sides for maxillary sinus) was used in the following analyses.

Only wild specimens were analyzed, as significant morphological differences were found between captive and wild specimens (Tables S6 and S7). Samples with missing values for sex (*N* = 1–2), hybrid index (*N* = 28), and hybrid class (*N* = 40–41) were excluded from subsequent analyses and figures requiring these variables. When comparing hybrid classes, the Oike individuals identified as P0 and the wild *M. cyclopis* from Izu-Oshima Island were treated together as one group (denoted as cyclopis_P0).

To evaluate the effect of hybridization on growth trajectories, the natural logarithm of centroid size (lnCS) was evaluated in relation to the developmental phase for each hybrid class. To summarize the shape variation of the cranium and mandible, PCA was performed on the shape components with all samples pooled using the “geomorph” R package, but each sex was shown separately in the scatterplots for better visualization. Differences in lnCS and PCs between hybrid classes at each stage were evaluated by analysis of variance with sex as a covariate using the “car” R package (Fox and Weisberg 2019), and their effect sizes were evaluated using the “rstatix” R package (Kassambara 2023).

Ontogenetic shape allometry was evaluated by multivariate regression using the R package “geomorph.” The response variable was the shape component, and the explanatory variables were lnCS, hybrid class, and their interaction. Pairwise vector correlations were evaluated using the R package “RRPP” (Collyer and Adams 2018, 2021). Phenotypic trajectory analysis was also performed to evaluate trajectory magnitude, direction, and shape differences between hybrid classes using the “geomorph” R package. Developmental stage, hybrid class, and their interaction were used as explanatory variables; hybrid classes with few samples in any of the three developmental stages (P0_cyclopis and F1 for females; F1 and F2 for males) were omitted. Disparities at each developmental stage were evaluated as a measure of Procrustes variance between the means of each hybrid class using the “geomorph” R package. For the maxillary sinus, reduced major axis regression was performed to assess the relationship between sinus lnCS and cranial lnCS, and slope and elevation differences between hybrid classes were tested using the “smatr” R package (Warton et al. 2012).

Adult variation was evaluated in relation to the hybrid index. To detect shape differences between pure *M. cyclopis* and *M. fuscata*, a between-group PCA (bgPCA) was performed, and admixed individuals were projected onto the bgPC1 axis using the “Morpho” R package (Schlager 2017). The lnCS, bgPC1, and relative sinus size (sinus lnCS – cranium lnCS) were regressed against the hybrid index for the admixed individuals, and deviations in elevation and slope from additivity (the line through the means of each parent) were tested using the “smatr” R package.

Deviations from additivity in multivariate space were evaluated as two orthogonal components, mismatch (*M*) and bias (*B*):

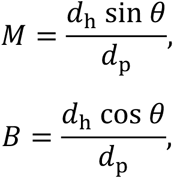

where *θ* is the angle between the vector passing through the mean of pure *M. cyclopis* and that of a hybrid class and the vector passing through the mean of pure *M. cyclopis* and that of *M. fuscata*; *d*_h_ and *d*_p_ are the magnitudes of the two vectors, respectively. For cranial and mandibular shapes, distances were calculated as Procrustes distances using the R package “shapes” (Dryden 2021). For relative maxillary sinus size, only *B* was calculated because it is univariate. *M* and *B* are similar to the transgression and closeness presented by Renaud et al. (2012), respectively, but differ in their orthogonality and are identical to the indices presented by Mérot et al. (2020) and Thompson et al. (2021), except for standardization. *M* > 0 indicates orthogonal deviations from the interparental axis, standardized by the interspecific distance. This is caused by different traits having dominance in conflicting directions in multivariate space (Thompson et al. 2021). Any deviations of *B* from the hybrid index indicate biases from additivity along the interparental axis, and in particular, *B* < 0 and *B* > 1 indicate exceeding the parental means, i.e., overdominance. *M* and *B* were calculated using a custom R script.

Each sex was analyzed separately in these analyses unless otherwise noted.

## 3. Results

### 3.1. Population structure

PCA and ADMIXTURE showed that the Oike population consisted of individuals with varying degrees of admixture, probably including pure *M. cyclopis* and *M. fuscata* (Figs. S1 and S2). The *M. cyclopis* individuals housed at the Primate Research Institute appeared to have a small amount of *M. fuscata*-derived ancestry (Fig. S2), but this did not affect subsequent analyses that included only samples from the Oike population. The NewHybrids analysis showed that the Oike population consists of the six predefined hybrid classes (P0, P1, F1, F2, BC0, and BC1; Fig. 3). It should be noted, however, that some of the individuals identified as F2, BC0, and BC1 may be from later generations. In the Oike population, the mean admixture proportion of *M. fuscata*-derived ancestry is 0.45 for autosomal, 0.46 for X-chromosomal in both sexes, and 0.50 for Y-chromosomal DNA. Note that the mean admixture proportion for autosomes was slightly biased towards *M. cyclopis* in BC0 and F2 than expected by random mating starting from the same number of parental lines (Table S8). The *M. fuscata*-type mtDNA ancestry was only observed in P0, i.e., pure *M. fuscata*. The P0 individuals were all males, while males were significantly less frequent than females in the F1 (*X*^2^ = 5.76, *Df* = 1, *P* = 0.016) (Table S9; Fig. 3). In the other hybrid classes, the sex ratio was not significantly skewed. The age structure was biased towards adults in F1 but towards earlier stages in F2 (Table S10; Fig. 3).

**Figure 3:**
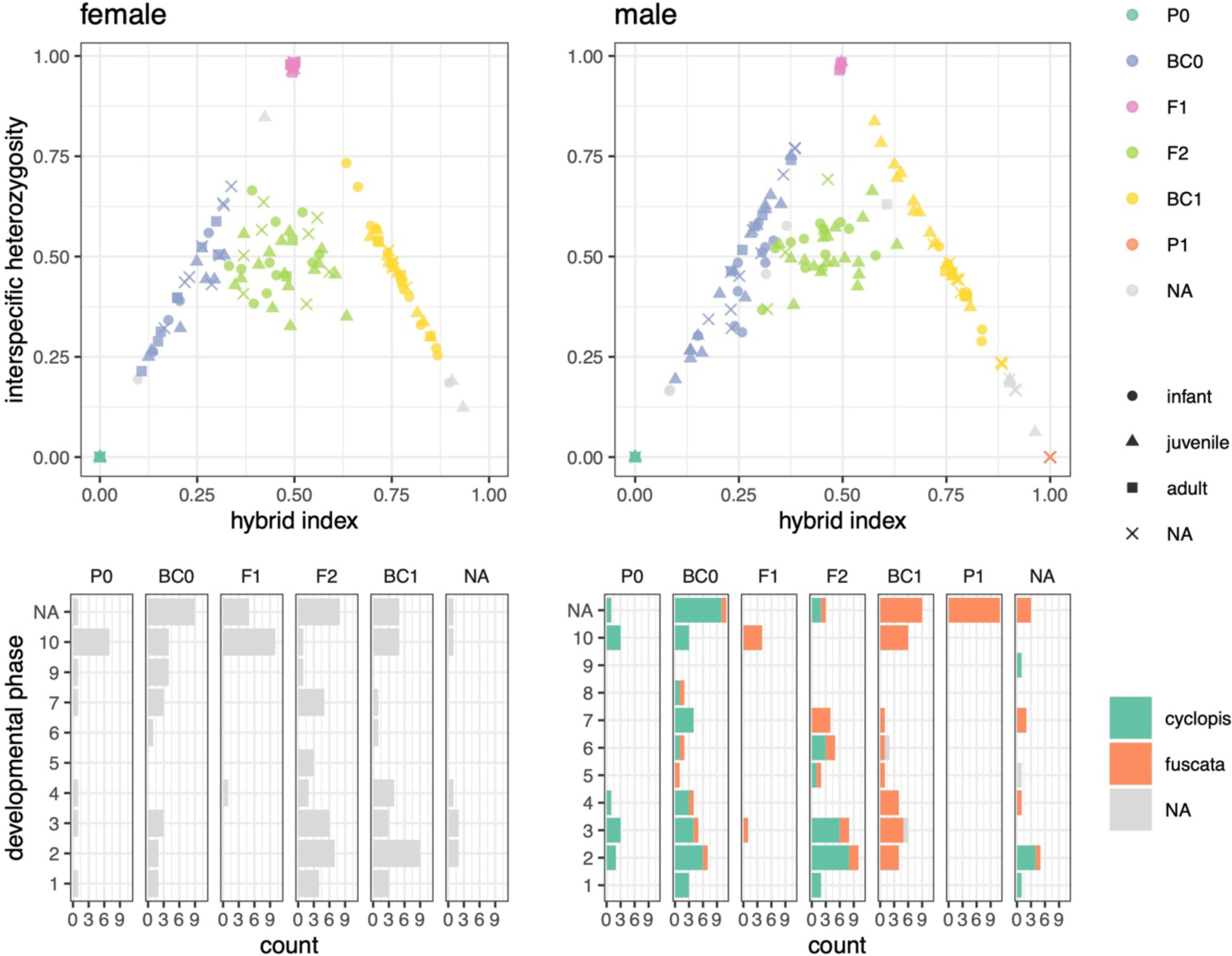
Population structure of the Oike population. The upper panels show the relationship between the hybrid index and interspecific heterozygosity with each colored by hybrid class. The lower panels show age structures; males are colored by Y chromosome ancestry.

### 3.2. Ontogenetic allometry

Throughout development, cranial and mandibular lnCS were larger in *M. fuscata* than in *M. cyclopis*, with hybrids in between in an almost additive manner (Table S11; Fig. S3). Among the first three PCs explaining >80% of the total variance, PC1 represented the ontogenetic allometric changes (Figs. 4 and S4), showing that larger (more developed) individuals had more elongated faces, relatively smaller neurocraniums, and narrower and longer mandibles. A similar shape change was observed in the multivariate regression analysis (see below). In contrast, PC2 represented interspecific differences throughout development (Table S11; Figs. 4, S4, and S5). *M. fuscata* showed a relatively longer face with an inferiorly inclined anterior maxilla and a mandible with a deeper ramus and shorter corpus than those of *M. cyclopis*; the hybrids were positioned in between in an almost additive manner. The PC plots indicated that the ontogenetic trajectories were not strictly linear, showing curves that detour in juveniles and return to an infantile shape (in the PC2 axis) in adults.

**Figure 4:**
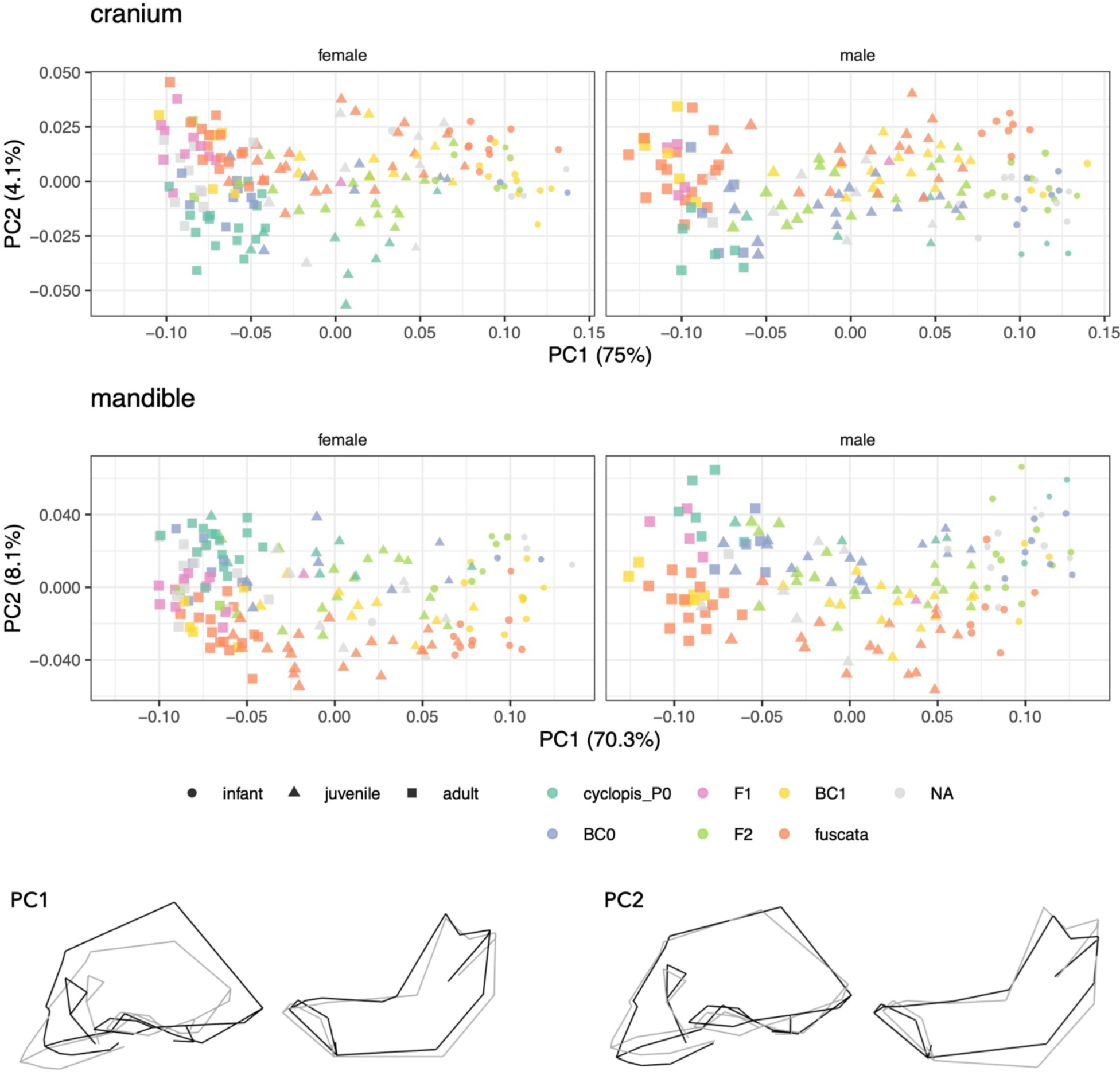
Variations of cranium and mandible shape represented by the first two principal components (PCs). All samples were analyzed together, but each sex was plotted separately for better visualization. Points are scaled according to lnCS. Shape changes along PC1 and PC2 are represented by wireframes (black, positive extreme; gray, negative).

Multivariate regression indicated that heterogeneity in allometric slopes was small to moderate but marginally significant in the female mandible and male cranium (Tables 1 and 2; Figs. 5 and S6). Allometric angles between the parental species ranged from 9.7 to 17.9 (degrees), although they were not significant after the Bonferroni adjustment. These values were comparable to those between a parental species and each of BC0, F2, and BC1 (8.3–18.7), although F1, which has few infant and juvenile specimens, often shows much larger angle differences (16.2–38.8). From infant to adult, the disparity decreased slightly in females and increased slightly in males (Fig. S7). The mismatch was already present in infants and/or juveniles to the extent of half the interspecific distance or less and remained constant or increased slightly until adulthood (Fig. S7). Allometric elevations were significantly different among hybrid classes, where hybrids were positioned between the parental species. Male allometric elevations were slightly biased towards *M. fuscata* than expected in the additive model, although the mean admixture proportion of autosomes was rather biased towards *M. cyclopis* in BC0 and F2 (Table S8). Phenotypic trajectory analysis supported the finding that ontogenetic trajectories were non-linear and did not differ greatly in magnitude, direction, and shape among hybrid classes, although in some cases, they were marginally significant (Table S12; Fig. 6).

**Figure 5:**
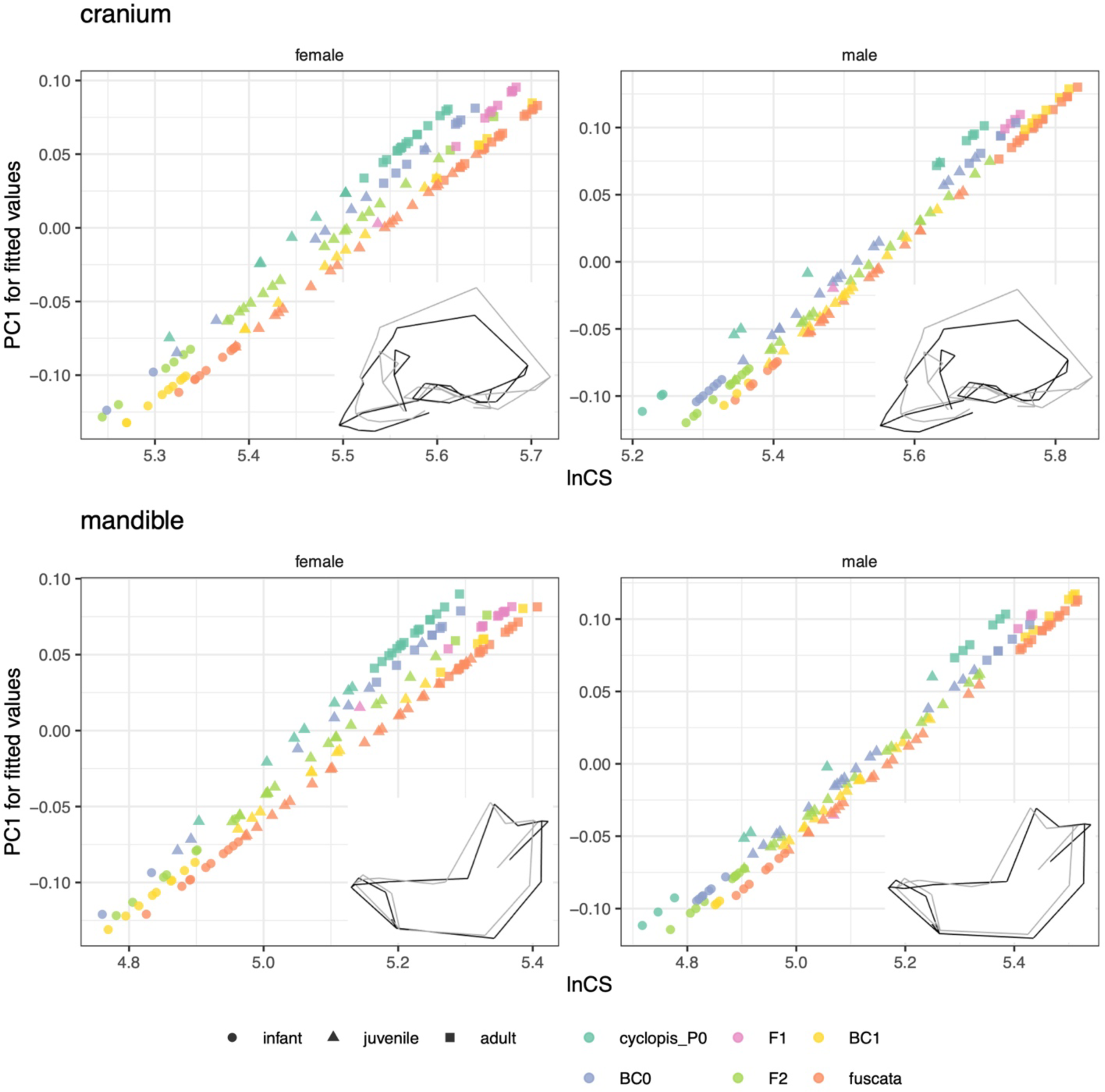
Ontogenetic allometry of cranial and mandibular shape represented by multivariate regression of shape against lnCS, hybrid class, and their interactions. Each sex was analyzed separately. Shape changes along PC1 for the fitted value are represented by wireframes (black, positive extreme; gray, negative).

**Figure 6:**
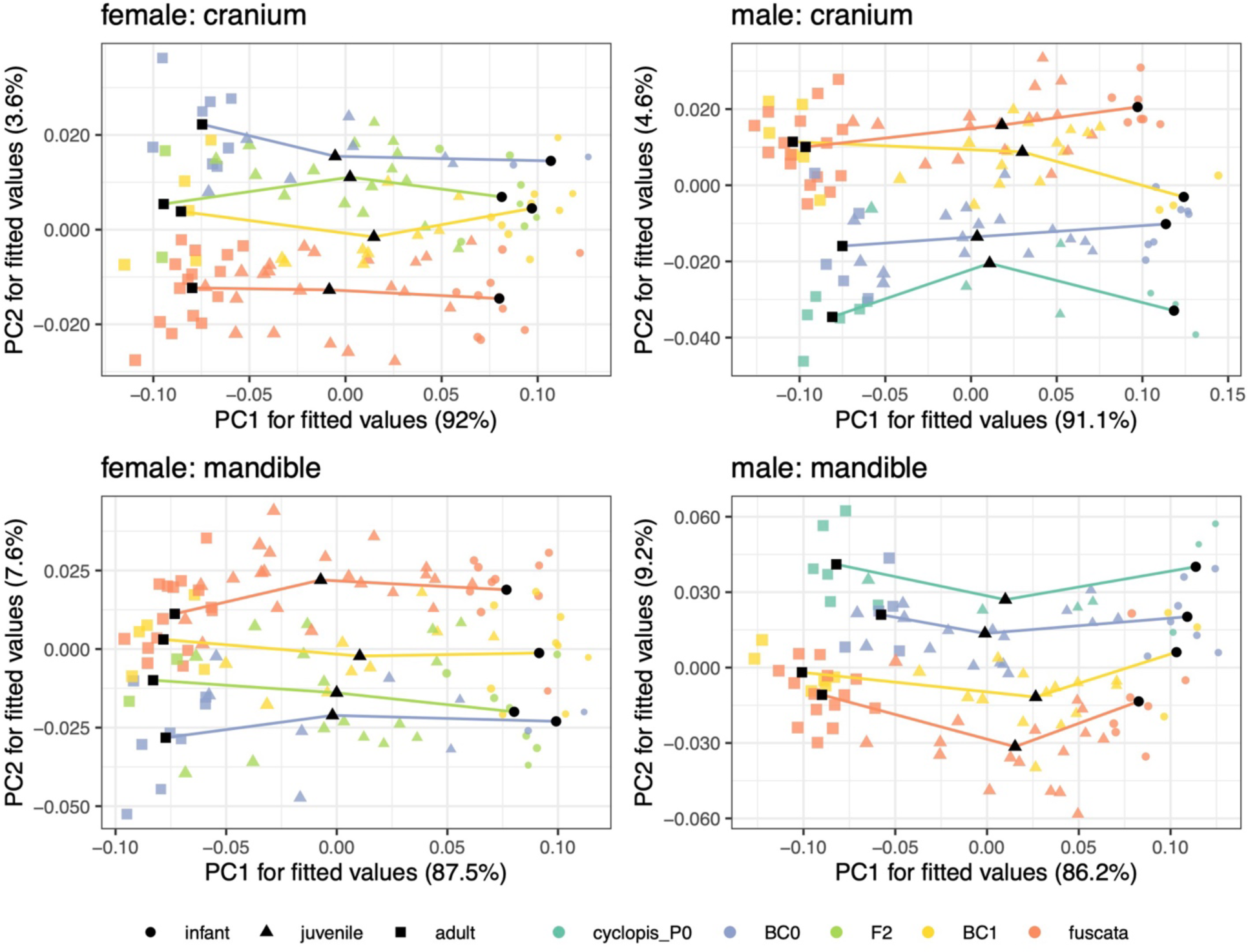
Ontogenetic trajectories of cranial and mandibular shape represented by the phenotypic trajectory analysis, where explanatory variables were lnCS, hybrid class, and their interactions. Each sex was analyzed separately. Hybrid classes with few samples (*M. cyclopis* and F1 for females; F1 and F2 for males) were excluded from the analyses. Points are scaled according to lnCS.

**Table 1.** Multivariate regression of shape on lnCS, hybrid class, and their interactions. Each sex was analyzed separately.

| Response | Source | *Df* | *SS* | *MS* | *R*<sup>2</sup> | *F* | *Z* | *P* |
| --- | --- | --- | --- | --- | --- | --- | --- | --- |
| *Female* |  |  |  |  |  |  |  |  |
| cranium | lnCS | 1 | 0.474 | 0.474 | 0.522 | 290.1 | 3.8 | **0.001** |
|  | hybrid class | 5 | 0.084 | 0.017 | 0.092 | 10.3 | 7.0 | **0.001** |
|  | lnCS:hybrid class | 5 | 0.010 | 0.002 | 0.011 | 1.2 | 1.4 | 0.074 |
|  | residuals | 134 | 0.219 | 0.002 | 0.241 |  |  |  |
|  | total | 145 | 0.909 |  |  |  |  |  |
| mandible | lnCS | 1 | 0.433 | 0.433 | 0.494 | 263.8 | 4.0 | **0.001** |
|  | hybrid class | 5 | 0.097 | 0.019 | 0.110 | 11.8 | 7.6 | **0.001** |
|  | lnCS:hybrid class | 5 | 0.012 | 0.002 | 0.013 | 1.4 | 2.2 | **0.015** |
|  | residuals | 133 | 0.218 | 0.002 | 0.249 |  |  |  |
|  | total | 144 | 0.877 |  |  |  |  |  |
| *Male* |  |  |  |  |  |  |  |  |
| cranium | lnCS | 1 | 0.691 | 0.691 | 0.659 | 414.9 | 3.4 | **0.001** |
|  | hybrid class | 5 | 0.059 | 0.012 | 0.056 | 7.1 | 7.6 | **0.001** |
|  | lnCS:hybrid class | 5 | 0.012 | 0.002 | 0.011 | 1.4 | 2.5 | **0.011** |
|  | residuals | 126 | 0.21 | 0.002 | 0.200 |  |  |  |
|  | total | 137 | 1.05 |  |  |  |  |  |
| mandible | lnCS | 1 | 0.582 | 0.582 | 0.614 | 319.9 | 3.7 | **0.001** |
|  | hybrid class | 5 | 0.067 | 0.013 | 0.070 | 7.3 | 8.0 | **0.001** |
|  | lnCS:hybrid class | 5 | 0.012 | 0.002 | 0.012 | 1.3 | 1.6 | 0.053 |
|  | residuals | 124 | 0.226 | 0.002 | 0.238 |  |  |  |
|  | total | 135 | 0.947 |  |  |  |  |  |

Type II sum of squares was used. Significance ( $P < 0.05$ ) is indicated in bold.

**Table 2.**
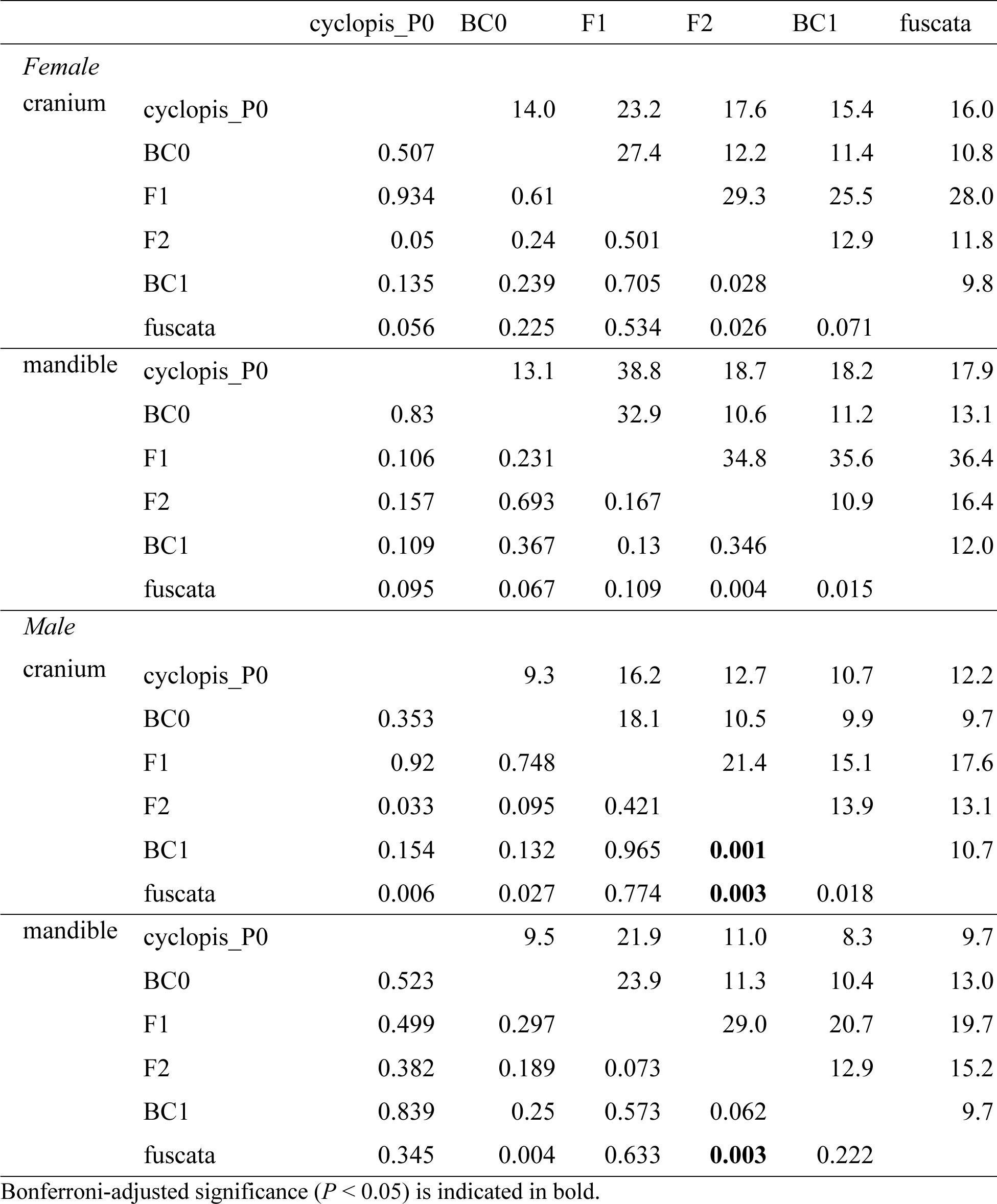
Pairwise vector correlations in the multivariate regression of shape against lnCS, hybrid class, and their interactions. The upper triangle indicates the angle (degrees), and the lower triangle indicates the *P* value.

The allometric slope of the maxillary sinus size differed significantly among the hybrid classes (Table S13; Fig. 7). There were small but significant differences in the absolute and relative sizes of the maxillary sinus, even at early developmental phases (Table S11; Fig. S3). *M. fuscata* had a milder allometric slope than the other hybrid classes, including *M. cyclops*, and the difference between *M. fuscata* and the others therefore increased at later developmental phases.

**Figure 7:**
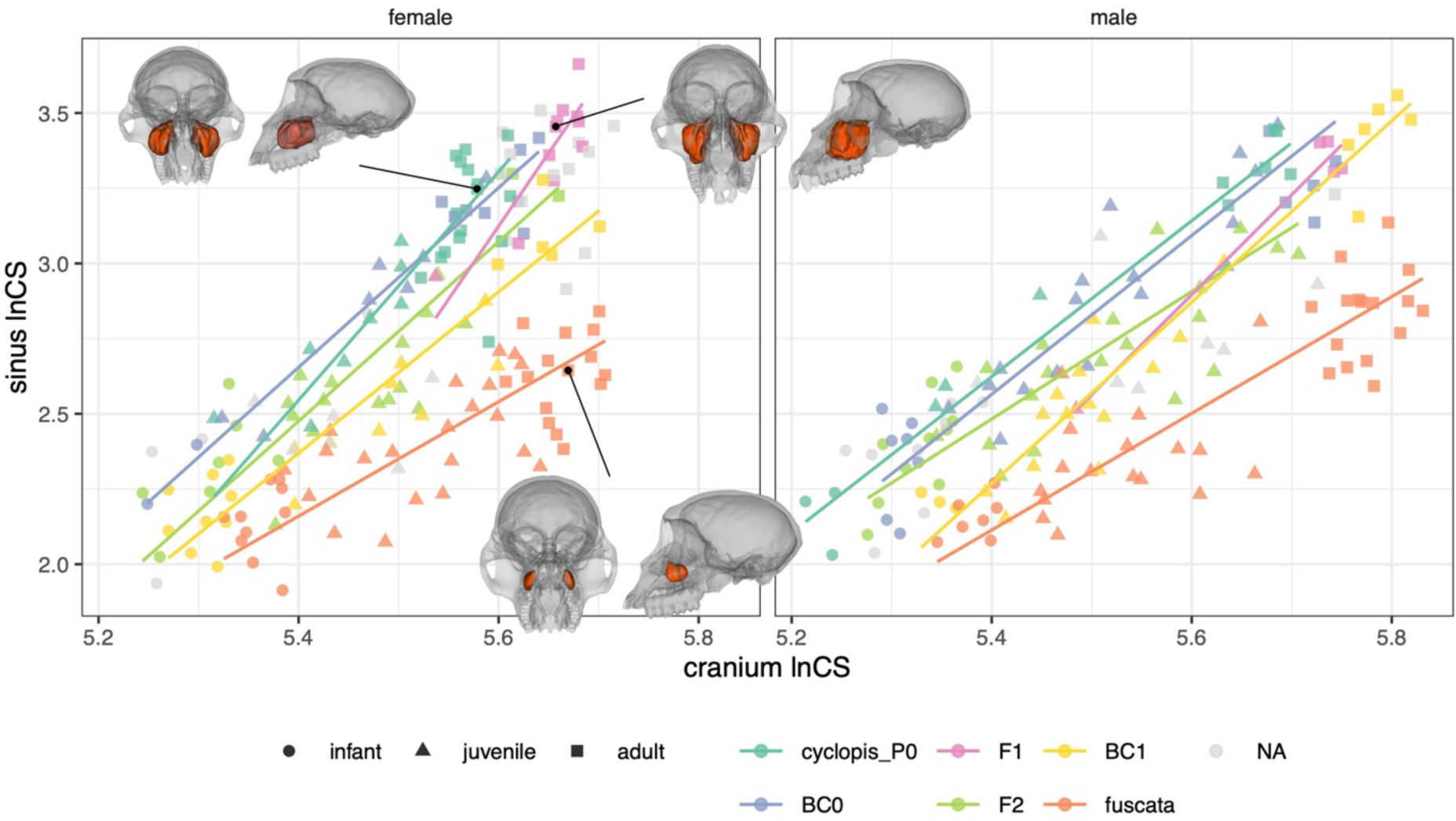
Ontogenetic allometry of maxillary sinus size versus the cranial size. The lines indicate the reduced major axis regression. The maxillary sinus is colored red in the transparent cranium.

### 3.3. Adult morphology

Admixed adults in the Oike populations followed the expectations of the additive model or were sometimes slightly biased towards *M. fuscata* in the size and shape component representing interspecific differences (bgPC1) (Table 3; Fig. 8). Differences in the major allometric components (PC1 and regression score) among hybrid classes were relatively small, although significant for cranial PC1 at developmental phase 10 (Table S11; Figs. 4, 5, and S6). In contrast, the maxillary sinuses of hybrids were as large as those of *M. cyclopis*, regardless of hybrid index, and deviated significantly from additivity (Table 3; Fig. 8). Marked transgressive segregations were observed in cranial and mandibular shapes (Table 4), particularly in the non-allometric shape space represented by PC2 and PC3 (Fig. S5). For cranial and mandibular shapes, deviations from additivity were relatively small in the bias component, while the degree of mismatch was comparable to half the interspecific distance. Some hybrids were also transgressive in the morpho-space of cranial shape that represents interspecific difference (PC2) and the relative size of the maxillary sinus (Fig. S8).

**Figure 8:**
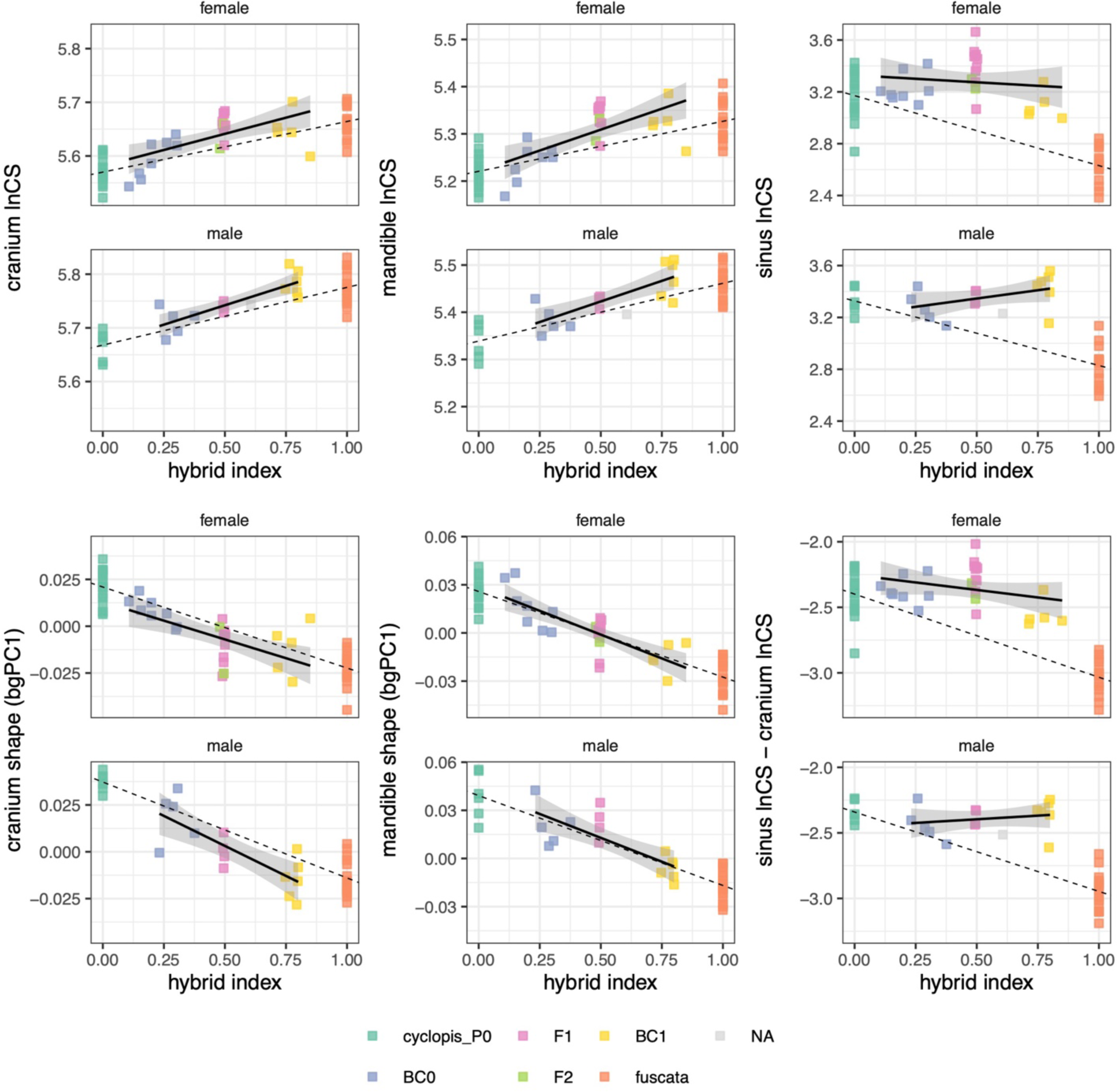
Relationship between morphological variation and hybrid index in adult samples. *M. fuscata* from the other populations and *M. cyclopis* are placed as 1 and 0 on the hybrid index, respectively. Dashed lines indicate the expectations of the additive model, which pass through the mean of the *M. cyclopis* (P0 and the other *M. cyclopis*) and the mean of *M. fuscata*. Solid lines indicate the ordinary least squares regression for the admixed individuals of the Oike population. For shape, between-group PCA (bgPCA) was performed to detect shape differences between pure *M. cyclopis* and *M. fuscata*, and admixed individuals were projected onto the bgPC1 axis.

**Table 3.**
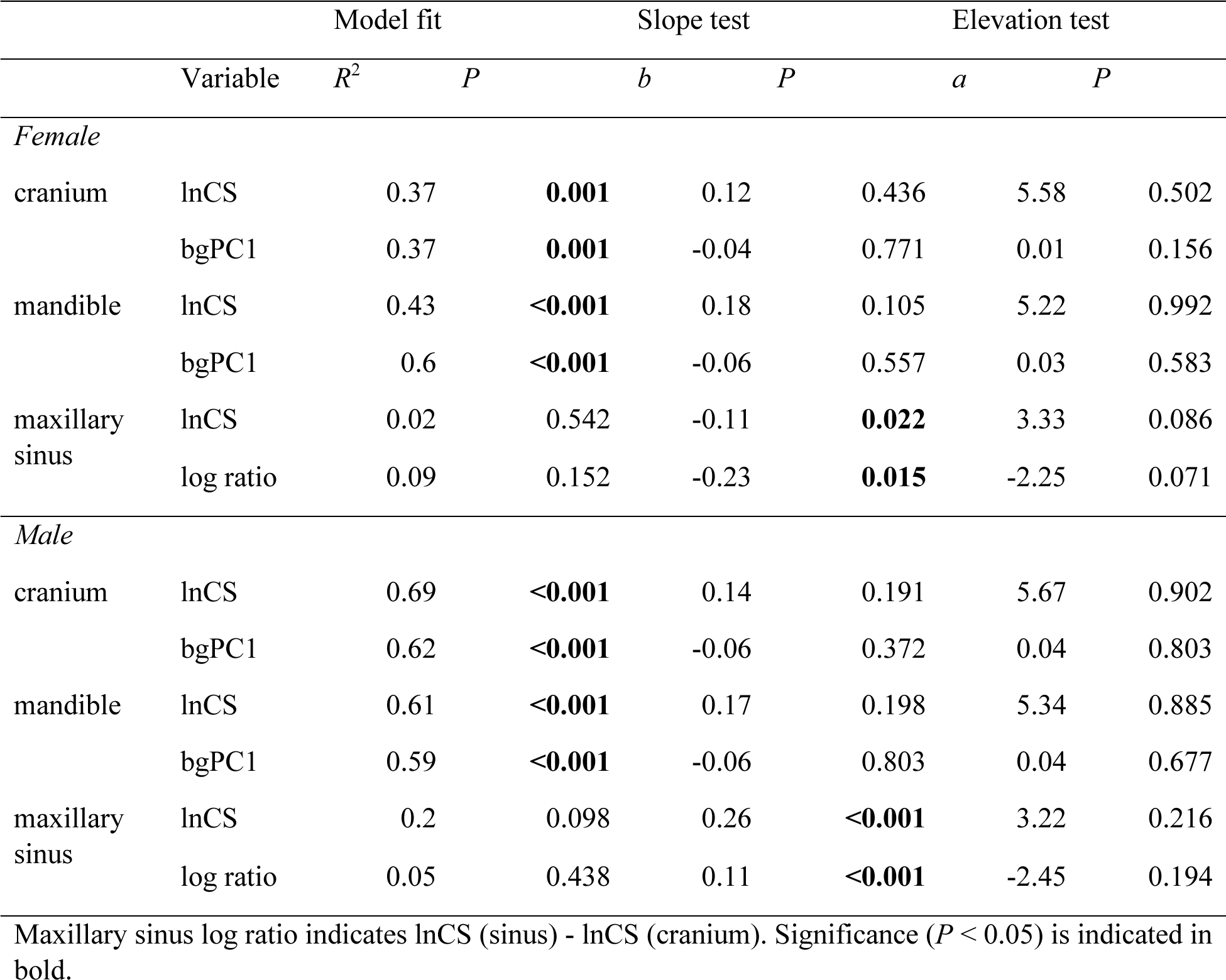
Regression of lnCS and bgPC1 on the hybrid index in the admixed adult samples. Deviations of elevation and slope from additivity (the line passing through the means of each parent) were evaluated. Each sex was analyzed separately.

**Table 4.** Transgressive segregation in multivariate shape space (but univariate size for maxillary sinus) represented by bias and mismatch in adult samples. A bias indicates the position of the data projected on the interspecific axis, where *M. cyclopis* is zero, and *M. fuscata* is one. A mismatch indicates orthogonal deviations from the interspecific axis standardized by the interspecific distance.

|  | BC0 |  | F1 |  | BC1 |  |
| --- | --- | --- | --- | --- | --- | --- |
|  | Female | Male | Female | Male | Female | Male |
| hybrid index | 0.21 | 0.29 | 0.50 | 0.49 | 0.77 | 0.78 |
| *Bias* |  |  |  |  |  |  |
| cranium | 0.30 | 0.36 | 0.74 | 0.72 | 0.77 | 1.01 |
| mandible | 0.18 | 0.33 | 0.51 | 0.30 | 0.77 | 0.81 |
| maxillary sinus | −0.05 | 0.16 | −0.24 | 0.07 | 0.24 | 0.03 |
| *Mismatch* |  |  |  |  |  |  |
| cranium | 0.56 | 0.48 | 0.82 | 0.60 | 0.53 | 0.55 |
| mandible | 0.34 | 0.62 | 0.58 | 0.49 | 0.53 | 0.51 |

## 4. Discussion

### 4.1. Population structure

This study examined 281 samples (after filtering) captured in the Oike area between 2003 and 2006. Considering that the estimated population size in the early 2000s was about 200–300 (Ohsawa et al. 2005) and the total number of captured individuals was 366, the samples used in this study were sufficient to evaluate the population structure of the Oike area.

Our results suggested that the Oike population consisted of pure *M. cyclopis*, pure *M. fuscata* (males only), the first and subsequent generations of hybrids, and the backcross hybrids of both parental species. The first generation consisted mainly of adults, while a few were observed in the subsequent generations. This implies that successful interbreeding between pure *M. cyclopis* and *M. fuscata* was probably sporadic and rare, at least in the years before eradication. In contrast, backcrossing and inter-hybrid breeding were probably widespread. As indicated in previous studies (Kawamoto et al. 2001, 2008a), no *M. fuscata*-type mtDNA was observed in the admixed individuals, suggesting that only male *M. fuscata* individuals migrated into the Oike population. Considering the philopatric nature of female macaques, this is a reasonable scenario. Given male migration and assuming random breeding, the proportion of Y-chromosome ancestry was expected to be twice that of autosomes, but this was much lower than expected. Therefore, a mechanism preventing random breeding, such as Haldane’s rule, population subdivision, or assortative mating, may have existed in this population. Fewer males than females in the F1 generation implies lower male survival or male emigration, although this is inconclusive due to the small sample size. The population subdivision scenario is also plausible, as four troops were observed in the Oike area (Ohsawa et al. 2005); heterogeneity in migration rate from neighboring *M. fuscata* between troops may underlie the relatively low frequency of *M. fuscat*a-type Y chromosomes in the Oike population.

### 4.2. Effects of hybridization on ontogenetic trajectories

In multivariate shape space, the ontogenetic trajectories of the cranium and mandible differed between the parental species. Direction differed (although not significantly) by less than 20 degrees between the parental species, which is comparable to that found between species within a papionin genus (Singleton 2012; Simons and Frost 2021). Hybridization shifted ontogenetic trajectories by the proportion of ancestry, although in some cases, with a slight bias toward *M. fuscata*. Hybridization also changed the direction to the same extent as the angle between the parental species (in most cases, they were not significant). However, the disparity and the degree of transgression did not greatly increase or decrease during development. In other words, the hybridization-induced shape differentiations were already manifested at birth; the differences in magnitude are maintained during postnatal ontogeny. This finding is partially consistent with a study of Sulawesi macaques, which showed no significant differentiation in the ontogenetic trajectory of the cranium between hybrids and their parental species in multivariate allometric space (Schillaci et al. 2005). Schillaci et al. (2005) reported hybridization-induced heterochrony in several cranial traits in bivariate allometric trajectories; this is pure heterochrony (*sensu* Mitteroecker et al. 2004), which cannot be evaluated when trajectories do not overlap in multivariate space. Thus, although hybridization shifts the ontogenetic trajectory and may even change the direction, these postnatal ontogenetic changes did not contribute to the increase in disparity and transgressive variation in adults. As suggested by Zelditch et al. (2016), the discrepancy may be ascribed to postnatally generated variation counteracting earlier generated variation.

It should be noted that the present finding of a limited contribution of hybridization-induced ontogenetic trajectory change to the increase in adult shape variation may be limited to the case of parental pairs with relatively small ontogenetic trajectory differences. Parallel ontogenetic trajectory differentiation between species has been widely observed in primate skull shape (Mouri 1996; Ponce de León and Zollikofer 2001; Toyoda et al. 2022), while recent multivariate studies have shown that the direction of ontogenetic trajectories of cranial shape differs between species to varying degrees across primate taxa (Mitteroecker et al. 2004; Vioarsdóttir and Cobb 2004; Singleton 2012; Simons and Frost 2021). The impacts of hybridization between species with greater differences in ontogenetic trajectories are of interest to future studies.

The finding that transgressive segregation is observed in hybrids is consistent with many previous studies examining hybrid phenotypic variation in various taxa of organisms (Rieseberg et al. 1999; Albertson and Kocher 2005; Stelkens et al. 2009; Renaud et al. 2012; Ackermann et al. 2019). What our finding can add to the literature is that transgressive segregation is likely to manifest prenatally without much change postnatally, at least in magnitude. Transgressive segregation is often found in a trait composed of multiple modules, such as the cranium and mandible examined in our study, and is probably caused by different modules responding to hybridization in opposite directions (Albertson and Kocher 2005; Renaud et al. 2012). Whether and how transgressive segregation manifests largely depends on the pattern and strength of integration among modules. Macaques appear to have achieved relaxed integration among modules in the cranium and mandible, respectively, to a degree sufficient to manifest transgressive segmentation.

The maxillary sinus of *M. cyclopis* is already larger than that of *M. fuscata* at birth, and the postnatal growth rate of *M. cyclopis* is much faster than that of *M. fuscata*. Hybridization altered the size of the maxillary sinus at birth by the proportion of ancestry, whereas the growth rate of hybrids was as fast as that of *M. cyclopis,* regardless of the proportion of ancestry. Although the genetic mechanisms underlying the non-additive inheritance of the postnatal growth rate of the maxillary sinus are unknown, two hypotheses can be proposed. First, *M. fuscata*-type cytoplasmic (e.g., mitochondrial) genes may inhibit sinus growth. Since cytoplasmic genes were not introduced into the hybrids due to male-biased migration, all hybrids exhibited a growth rate as fast as that of *M. cyclopis*. Maternal inheritance associated with nuclear genes, such as genomic imprinting, would not account for the deviations in the second and subsequent generations of hybrids. Second, an epistatic interaction of *M. fuscata*-type alleles may inhibit sinus growth. Such an interaction can be disrupted by the introgression of alleles from another species, even if only a small fraction. Considering that backcrossed individuals did not recover the slow sinus growth rate typical of *M. fuscata*, an epistatic interaction, if any, was controlled by not a few genes. In either case, as a consequence, the adult hybrids show a mosaic pattern, i.e., the maxillary sinus is as large as that of *M. cyclopis*, although the cranium shape is almost intermediate (in the shape component representing interspecific difference) between the two parental species.

### 4.3. Evolutionary background underlying the consequences of hybridization

How hybridization affects ontogenetic trajectories is likely to depend on the evolutionary processes that produced the difference in genetic architecture between the parental lineages. This idea stems from the finding that genetic architecture imposes limits on transgressive segregation (Albertson and Kocher 2005). The whole body shape of fish shows much greater differences in ontogenetic direction between the parental lineages and between them and their hybrids (Sinama et al. 2013; Santos-Santos et al. 2021) than those observed in the cranium and mandible of hybrid macaques. It can be hypothesized that the fish body is less integrated between modules and has experienced selective pressures that act differently on each module, producing greater changes in ontogenetic direction. In contrast, the evolutionary forces shaping interspecific differences in macaque skulls may not have been strong enough to greatly affect ontogenetic directions.

The genetic distance between the parental lines is a significant predictor of the expression of transgressive segregation in hybrids (Stelkens and Seehausen 2009; Stelkens et al. 2009). The evolutionary history of the group consisting of *M. cyclopis*, *M. fuscata*, and *M. mulatta* has been relatively well studied. Among the three, *M. fuscata* diverged first, followed by *M. cyclopis*, and then the Chinese and Indian lineages of *M. mulatta* diverged, with gene flow between some pairs of their ancestors (Osada et al. 2021). Although the time of divergence between *M. cyclopis* and *M. fuscata* remains controversial, the two species were probably isolated no later than 0.43 million years ago, the last time the Japanese archipelago was connected to the continent (Aimi 2002). It should be noted that the substitution rate of the *M. fuscata* lineage is much higher than the other lineages, probably due to the ancestral population bottleneck and the resulting accumulation of slightly deleterious mutations (Ito et al. 2021). *M. fuscata* is also peculiar in that it has the northernmost distribution among nonhuman primates, and some of its morphological features probably reflect adaptation to high-latitude environments (Antón 1996). Such relatively deep divergence and specialization in the parental lineages probably formed the genetic architecture that produced transgressive phenotypes in hybrids. This is in contrast to the case of pelvic morphology of hybrids between Chinese and Indian lineages of *M. mulatta*, which shows weak inter-lineage differences and no evidence of transgressive phenotypes (Buck et al. 2021). The divergence between Chinese and Indian lineages of *M. mulatta* is shallower than that between *M. cyclopis* and *M. mulatta*, but not as different. The demographic history and adaptive evolution, not just the depth of divergence, of the parental lineages is likely to be the key to what variation is manifested in hybrids.

The maxillary sinus is peculiar in that the growth rate appears to show cytoplasmic or epistatic inheritance. If epistatic interaction inhibits sinus growth in *M. fuscata*, it is reasonable to assume that the genetic architecture has been formed by natural selection, i.e., a coadapted gene complex. The small maxillary sinus may be advantageous in the high-latitude environment inhabited by *M. fuscata* because it could provide space for the expansion of the nasal cavity or to support a strong bite. In fact, *M. fuscata* populations at higher latitudes have a larger nasal cavity and a smaller maxillary sinus (Rae et al. 2003), and *M. fuscata* shows a cranial adaptation to a high attrition diet compared to its relatives (Antón 1996). Alternatively, the growth rate of the maxillary sinus may be controlled by a genetic factor that pleiotropically affects an unknown functional trait, as it is questionable whether the maxillary sinus has a functional role (Rae and Koppe 2004; Márquez 2008) or is related to biomechanical force (Rae and Koppe 2008). In either case, a somewhat independent trait that is under different selective pressure from the surrounding structure, such as the maxillary sinus, may lead to specific ontogenetic trajectory differentiation in hybrids, whereas highly integrated traits may not.

### 4.3. Conclusions

This study showed that: (1) Hybridization shifted the ontogenetic trajectories of cranial and mandibular shapes in a nearly additive manner. (2) Transgressive variation in cranial and mandibular shape manifested at birth; hybridization-induced changes in postnatal ontogenetic direction did not contribute to increases in adult transgressive variation or disparity. (3) Hybridization disrupted a *M. fuscata*-specific genetic function that inhibited maxillary sinus growth, producing hybrids with a mosaic pattern, i.e., the maxillary sinus was as large as that of *M. cyclopis*, although the cranial shape was intermediate in the shape component representing interspecific difference. This study has several limitations. First, the skeletal specimens used as surrogates for the parental populations may differ in morphology from the actual parental populations due to regional intraspecific variation, although the surrogates were selected from localities as close as possible to the locality of the hybrid population. Second, the ontogenetic trajectories lack stability due to the small sample size, and the present findings should be validated based on a larger sample size. Third, the F2, BC0, and BC1 categorized in this study may include later generations, and such genetic heterogeneity of group members may obscure patterns of variation; this concern could be addressed by inferring generations in detail based on fine-scale phased data or using the samples with known genealogy. Despite these limitations, this study highlights the complex genetic and ontogenetic bases underlying hybridization-induced differentiation in ontogenetic trajectories and resulting morphological diversification.

## Author Contributions

TI, RK, YH, and YK conceived the research. TI, HW, and YK performed basic molecular experiments. AT and AJN conducted ddRAD library preparation. YK contributed samples and data. TI and MT assessed the dental eruption-cranial suture stage. TI analyzed the data and drafted the manuscript. All authors approved the final version of this manuscript.

## Acknowledgments

We thank Hideyuki Ohsawa, Yoshiki Morimitsu, Yasuyuki Muroyama, Shingo Maekawa, Hideo Nigi, Harumi Torii, Shunji Goto, Tamaki Maruhashi, Naofumi Nakagawa, Jun Nakatani, Toshiki Tanaka, Sachiko Hayakawa, Aya Yamada, Shuhei Hayaishi, Hironori Seino, Kei Shirai, Mami Saeki, Shizuka Kawai, Ko Hagiwara, Katsuya Suzuki, Kunihiko Suzuki, Junya Uetsuki, Misao Okano, Tadanobu Okumura, Atsuhisa Yoshida, and Noriko Yokoyama, of the Working Group of Wakayama Taiwanese macaque, and the Department of Environment and Life, Wakayama Prefecture, for kindly allowing us to use the samples. We also thank Yuta Shintaku and Takeshi Nishimura for their help in accessing the skeletal collections. We are grateful to Akira Kawaguchi, Kyoko Yamaguchi, Takehiro Sato, and Chikatoshi Sugimoto for teaching molecular experiments to TI. We express our gratitude to the successive members of the Department of Evolution and Phylogeny, Primate Research Institute, Kyoto University, for their efforts in collecting, preparing, and preserving skeletal collections. This study was funded by the Keihanshin Consortium for Fostering the Next Generation of Global Leaders in Research (K-CONNEX) established by the Human Resource Development Program for Science and Technology, MEXT, JSPS KAKENHI (Grant Numbers: JP15J00134, JP17K15195, JP19K16211, and JP19H01002), the University of the Ryukyus Research Project Program Grant for Junior Researchers, the Cooperative Research Program from the Primate Research Institute of Kyoto University, and ISHIZUE 2022 of Kyoto University o TI.

## Data Accessibility Statement

The sequence reads have been submitted to the Sequence Read Archive (SRA) of the National Center for Biotechnology Information (NCBI) under accession number PRJNA1009132. The other data underlying this article are available in the Dryad Digital Repository, at https://doi.org/10.5061/dryad.d2547d87h. (These data will be available upon acceptance of this article.)

