## Supplementary materials for "Hybridization and its impact on the ontogenetic allometry of skulls in macaques"

### Supplementary Note

#### S1. Double-digest restriction site-associated DNA library preparation and sequencing

Genomic DNA (10 ng) was digested with EcoRI and BglII (New England Biolabs, Ipswich, MA, USA), and adapters were ligated at 37 °C overnight in a 10-μL volume containing 1 μL of ×10 NEB buffer 2, 0.1 μL of ×100 bovine serum albumin (New England Biolabs), 0.4 μL of 5 μM EcoRI adapter and BglII adapter, 0.1 μL of 100 mM ATP and 0.5 μL of T4 DNA Ligase (Enzymatics, Beverly, MA, USA). The reaction solution was purified with AMPure XP (Beckman Coulter). Next, 3 μL of purified DNA was used in the PCR amplification in a 10-μL volume containing 1 μL of each 10 μM index and TruSeq universal primer, 0.3 μL of KOD-Plus-Neo enzyme, and 1 μL of ×10 PCR buffer (Toyobo, Osaka, Japan), 0.6 μL of 25 mM MgSO4 and 1 μL of 10 mM dNTP. Thermal cycling was initiated with a 94 °C step for 2 min, followed by 20 cycles of 98 °C for 10 s, 65 °C for 30 s, and 68 °C for 30 s. PCR products were pooled and purified again with AMPure XP. Purified DNA was loaded onto a 2.0% agarose gel, and fragments of around 360 base pairs (bp) were retrieved using E-Gel SizeSelect (Life Technologies, Carlsbad, CA, USA). After quality assessment using an Agilent 2100 Bioanalyzer (Agilent Technologies, Santa Clara, CA, USA), the library was sequenced with 51-bp single-end reads in six lanes of an Illumina HiSeq2000 (Illumina, San Diego, CA, USA) by Macrogen Inc. (Seoul, South Korea). Primer sequences used in this study are available in Sakaguchi et al. (2015).

#### S2. Variant calling

Raw reads were filtered for overall sequence quality using TRIMMOMATIC 0.39 (Bolger et al. 2014) with the following parameters: ILLUMINACLIP:TruSeq3-SE.fa:2:30:10, LEADING:19,

TRAILING:19, SLIDINGWINDOW:30:20, and AVGQUAL:20 MINLEN:51. Filtered reads were mapped to the *M. mulatta* (rhesus macaque) RefSeq (Mmul_10; GCF_003339765.1) using BOWTIE2 2.4.4 (Langmead and Salzberg 2012) with --very-sensitive option. The sex of each individual was assessed based on the ratio of the number of reads mapped to the Y chromosome to those mapped to the X chromosome using sam_sexing.py (https://github.com/itots/sam_sexing.git). Genotypes were called using STACKS 2.59 (Rochette and Catchen 2017), using the Marukilow model in the gstacks function to search for variant sites with relatively stricter criteria than the default setting: --var-alpha 0.01 (a significant level for calling variant sites), --gt-alpha 0.01 (a significant level for calling genotypes), and --min-mapq 20 (minimum PHRED-scaled mapping quality to consider a read). Next, the populations function was used to call SNPs with the following parameters: --rite-single-snp (extract only the first SNP per locus) and --ordered-export.

**S3. Variant filtration**

The SNPs dataset was filtered using PLINK 1.90b6.21 (Purcell et al. 2007; https://www.cog-genomics.org/plink2/) to retain individuals with missing rate <0.2 and sites with missing rate <0.2 and minor allele frequency >0.01 (--geno 0.2 --mind 0.2 --maf 0.01). Autosomal and X chromosome SNPs were further filtered to remove sites in strong linkage disequilibrium (--indep-pairwise 10 3 0.5). These filtrations yielded 6461 variant sites in 289 individuals for the autosome (total genotyping rate = 0.96), 171 sites in 289 individuals for the X chromosome (0.96), and 12 sites in 152 individuals for the Y chromosome (0.95) DNA.

**S4. Data screening**

Data from Kawamoto et al. (2008a) were also examined, including sex and a nine-step hybrid index calculated from four autosomal markers (blood proteins of transferrin, adenosine deaminase, diaphorase, and SNP of natural resistance-associated macrophage protein 1) and ancestry types of mitochondrial DNA (mtDNA) and testis-specific protein Y (TSPY). Some missing data on Y chromosome ($N = 52$) and mtDNA ancestry ($N = 9$) were supplemented in this study; the method used was identical to that of Kawamoto et al. (2008a), but when the results of the TSPY marker were unclear, two STR markers (DYS472 and DYS569) were additionally examined according to Kawamoto et al. (2008b).

Data from Kawamoto et al. (2008a) and this study were compared to screen the data. Sex identification was inconsistent in five samples. The ADMIXTURE $Q$ score ($K = 2$) was highly correlated with the nine-step hybrid index ($R^2 = 0.79$, $F_{1, 275} = 1005.7$, $P < 0.001$), but two samples were detected as outliers in their residual component (Bonferroni adjusted $P < 0.05$). These six samples (one sample was inconsistent in both tests) were removed from the analyses. Y-chromosome ancestry was identical between the two datasets, except for three samples with missing values in Kawamoto et al. (2008a).

**S5. 3D data acquisition**

Craniums and mandibles were scanned by computed tomography (CT) at the Primate Research Institute using the Asteion Premium 4 helical scanner (Toshiba Medical Systems, Otawara, Japan) with a slice thickness of 0.5 mm. Serial CT images were reconstructed from the original volumetric data with a pixel size between 0.093 and 0.314 mm and an interslice interval between 0.1 and 0.3 mm, depending on the specimen size. CT images were segmented based on edge detection and watershed methods (Ito 2019), and three-dimensional (3D) surface models were created using Python (create_mesh.py; https://github.com/itots/dicom2stl.git). Image stacks and the 3D models were loaded into 3D Slicer (https://www.slicer.org/) (Kikinis et al. 2014) for landmark digitization.

Purcell, S., B. Neale, K. Todd-Brown, L. Thomas, M. A. R. Ferreira, D. Bender, J. Maller, P. Sklar, P. I.

W. de Bakker, M. J. Daly, and P. C. Sham. 2007. PLINK: A tool set for whole-genome association and population-based linkage analyses. Am. J. Hum. Genet. 81:559–575.

R Developmental Core Team. 2021. R: A language and environment for statistical computing. Vienna.

Rochette, N. C., and J. M. Catchen. 2017. Deriving genotypes from RAD-seq short-read data using Stacks. Nat. Protoc. 12:2640–2659.

Sakaguchi, S., K. Horie, N. Ishikawa, A. J. Nagano, M. Yasugi, H. Kudoh, and M. Ito. 2017. Simultaneous evaluation of the effects of geographic, environmental and temporal isolation in ecotypic populations of Solidago virgaurea. New Phytol. 216:1268–1280.

Sakaguchi, S., T. Sugino, Y. Tsumura, M. Ito, M. D. Crisp, D. M. J. S. Bowman, A. J. Nagano, M. N. Honjo, M. Yasugi, H. Kudoh, Y. Matsuki, Y. Suyama, and Y. Isagi. 2015. High-throughput linkage mapping of Australian white cypress pine (*Callitris glaucophylla*) and map transferability to related species. Tree Genet. Genomes 11:1–12.

#### Supplementary Tables

Table S1: Total samples used in this study. The developmental (dev.) stage was determined based on the developmental phase. Note that only wild samples were used for the main analyses.

| | | | | Skull | | | DNA | |
|---|---|---|---|---|---|---|---|---|
| Source | Environment | Locality | Dev. stage | Female | Male | Unknown | Female | Male |
| Oike population | wild | Oike | infant | 35 | 37 | | 30 | 34 |
| | | | juvenile | 44 | 62 | 2 | 39 | 59 |
| | | | adult | 45 | 20 | | 34 | 19 |
| | | | unknown | | | | 33 | 39 |
| *M. cyclopis* (ref.) | captive | unknown | infant | 2 | 3 | | | |
| | | | juvenile | 7 | 5 | 1 | 1 | |
| | | | adult | 2 | 3 | | | 1 |
| | | | unknown | | | | 1 | 1 |
| | wild | Izu-Oshima | infant | | 1 | 1 | | |
| | | | juvenile | 5 | 1 | 1 | | |
| | | | adult | 8 | 3 | | | |
| | unknown | unknown | infant | 1 | | | | |
| | | | juvenile | 3 | | 1 | | |
| | | | adult | 2 | 4 | | | |
| *M. fuscata* (ref.) | captive | Kyoto | juvenile | | 1 | | | 1 |
| | | | adult | | 1 | | | 1 |
| | | | unknown | | | | 2 | |
| | wild | Noguchi | adult | | 1 | | | |
| | | Shiga | infant | 7 | 5 | | | |
| | | | juvenile | 15 | 10 | | | |
| | | | adult | 7 | 5 | | | |
| | | Fukui | juvenile | 2 | 1 | | | |
| | | | adult | 5 | 7 | | | |
| | | Shimane | infant | 3 | 2 | | | |
| | | | juvenile | 4 | 6 | | | |
| | | | adult | 3 | 3 | | | |

Table S2: Definition of developmental (dev.) stage and phase.

| Dev. stage | Dev. phase | Definition |
|---|---|---|
| infant | 1 | Before dental stage 2 |
| | 2 | The full eruption of all the deciduous teeth |
| juvenile | 3 | The full eruption of all the four permanent first molars |
| | 4 | The full eruption of all the four permanent first incisors |
| | 5 | The full eruption of all the four permanent second incisors |
| | 6 | The full eruption of all the four permanent second molars |
| | 7 | The full eruption of all the eight permanent premolars |
| adult | 8 | The full eruption of all the four permanent third molars, but before the full eruption of the four permanent canines |
| | 9 | The full eruption of all the permanent teeth, but before the complete closure of spheno-occipital synchondrosis |
| | 10 | The full eruption of all the permanent teeth and the complete closure of spheno-occipital synchondrosis |

Table S3: Landmarks used in this study.

| No | | | | |
|---|---|---|---|---|
| Middle | Left | Right | Landmarks | Module |
| **Cranium** | | | | |
| 1 | | | Prosthion | Face |
| 2 | | | Nasospinale | Face |
| 3 | | | Rhinion | Face |
| 4 | | | Nasion | Face |
| 5 | | | Glabella | Brain |
| 6 | | | Bregma | Brain |
| 7 | | | Inion | Brain |
| 8 | | | Opisthion | Brain |
| 9 | | | Basion | Brain |
| 10 | | | Hormion | Brain |
| 11 | | | Staphylion | Face |
| 12 | | | Incisivision | Face |
| | 13 | 32 | Distal I2 (i2) (lateral) | Face |
| | 14 | 33 | Distal C (c) (lateral) | Face |
| | 15 | 34 | Distal P4 (dm2) (lateral) | Face |
| | 16 | 35 | Maxilla-palatine (inferior) | Face |
| | 17 | 36 | Nasal–premaxilla | Face |
| | 18 | 37 | Dacryon | Face |
| | 19 | 38 | Zygo-max superior | Face |
| | 20 | 39 | Zygo-max inferior | Face |
| | 21 | 40 | Frontomalare orbitale | Face |
| | 22 | 41 | Jugale | Face |
| | 23 | 42 | Zygo-temp inferior | Face |
| | 24 | 43 | Zygomatic arch–alishpenoid (inferior) | Brain |
| | 25 | 44 | Sphenoasquamosal suture on infratemporal crest | Brain |
| | 26 | 45 | Temporal zygomatic curve (posterior) | Brain |
| | 27 | 46 | Porion | Brain |
| | 28 | 47 | Asteion | Brain |
| | 29 | 48 | Jugular foramen (lateral) | Brain |
| | 30 | 49 | Carotid canal (anterior) | Brain |
| | 31 | 50 | Occipital condyle (posterior) | Brain |
| **Mandible** | | | | |
| 1 | | | Infradentale | |
| 2 | | | Mandibular orale | |
| 3 | | | Superior transverse torus (posterior) | |
| 4 | | | Gnathion | |
| | 5 | 17 | Mental foramen (anterior) | |

Table S3: Landmarks used in this study. *(continued)*

| No | | | | |
|---|---|---|---|---|
| Middle | Left | Right | Landmarks | Module |
| | 6 | 18 | Distal I2 (i2) (lateral) | |
| | 7 | 19 | Distal C (c) (lateral) | |
| | 8 | 20 | Distal P4 (dm2) (lateral) | |
| | 9 | 21 | Ramus (anterior and in line with alveolus) | |
| | 10 | 22 | Gonion | |
| | 11 | 23 | Ramus (posterior and in line with alveolus) | |
| | 12 | 24 | Condylion laterale | |
| | 13 | 25 | Condylion mediale | |
| | 14 | 26 | Sigmoid notch | |
| | 15 | 27 | Coronion | |
| | 16 | 28 | Mandibular foramen (superior) | |
| **Maxillary sinus** | | | | |
| | 1 | 2 | Anterior | |
| | 3 | 4 | Posterior | |
| | 5 | 6 | Inferior | |
| | 7 | 8 | Lateral | |
| | 9 | 10 | Superior | |

Table S4: Measurement errors of shape evaluated by Procrustes analysis of variance.

| Source | $Df$ | $SS$ | $MS$ | $R^2$ | $F$ | $Z$ | $P$ |
|---|---|---|---|---|---|---|---|
| **Cranium** | | | | | | | |
| ind | 355 | 10.49 | 0.0295 | 0.953 | 39.7 | 11.6 | **0.001** |
| side | 1 | 0.05 | 0.0478 | 0.004 | 64.3 | 10.0 | **0.001** |
| ind:side | 355 | 0.26 | 0.0007 | 0.024 | 2.5 | 29.6 | **0.001** |
| ind:side:replicate | 712 | 0.21 | 0.0003 | 0.019 | | | |
| total | 1423 | 11.00 | | | | | |
| **Mandible** | | | | | | | |
| ind | 342 | 9.66 | 0.0282 | 0.954 | 35.0 | 8.3 | **0.001** |
| side | 1 | 0.03 | 0.0294 | 0.003 | 36.4 | 8.7 | **0.001** |
| ind:side | 342 | 0.28 | 0.0008 | 0.027 | 3.5 | 27.2 | **0.001** |
| ind:side:replicate | 686 | 0.16 | 0.0002 | 0.016 | | | |
| total | 1371 | 10.12 | | | | | |
| **Maxillary sinus** | | | | | | | |
| ind | 358 | 53.12 | 0.1484 | 0.810 | 7.2 | 7.4 | **0.001** |
| side | 1 | 0.15 | 0.1495 | 0.002 | 7.2 | 4.4 | **0.001** |
| ind:side | 358 | 7.42 | 0.0207 | 0.113 | 3.0 | 23.4 | **0.001** |
| ind:side:replicate | 718 | 4.91 | 0.0068 | 0.075 | | | |
| total | 1435 | 65.60 | | | | | |

*Note:*
Type I sum of squares is used. Significance (< 0.05) is indicated in bold.

Table S5: Measurement errors of centroid size evaluated by analysis of variance.

| Source | $Df$ | $SS$ | $MS$ | $F$ | $P$ |
| --- | --- | --- | --- | --- | --- |
| **Cranium** | | | | | |
| ind | 355 | 988397.2 | 2784.22 | 17970.8 | **< 0.001** |
| residuals | 356 | 55.2 | 0.15 | | |
| **Mandible** | | | | | |
| ind | 342 | 797735.5 | 2332.56 | 44514.4 | **< 0.001** |
| residuals | 343 | 18.0 | 0.05 | | |
| **Maxillary sinus** | | | | | |
| ind | 358 | 78787.8 | 220.08 | 447.2 | **< 0.001** |
| residuals | 1077 | 530.1 | 0.49 | | |

*Note:*
Significance (< 0.05) is indicated in bold.

Table S6: Effect of the environment (captive–wild difference) on shape evaluated by Procrustes analysis of variance. Developmental phase, species, and sex are used as covariates.

| Source | $Df$ | $SS$ | $MS$ | $R^2$ | $F$ | $Z$ | $P$ |
|---|---|---|---|---|---|---|---|
| **Cranium** | | | | | | | |
| Developmental phase | 1 | 0.48 | 0.48 | 0.60 | 240.3 | 3.9 | **0.001** |
| species | 1 | 0.04 | 0.04 | 0.04 | 17.5 | 5.4 | **0.001** |
| sex | 1 | 0.01 | 0.01 | 0.01 | 5.0 | 3.5 | **0.001** |
| environment | 1 | 0.01 | 0.01 | 0.01 | 4.1 | 3.2 | **0.001** |
| residuals | 119 | 0.24 | 0.00 | 0.30 | | | |
| total | 123 | 0.81 | | | | | |
| **Mandible** | | | | | | | |
| Developmental phase | 1 | 0.42 | 0.42 | 0.55 | 191.9 | 4.1 | **0.001** |
| species | 1 | 0.04 | 0.04 | 0.06 | 19.5 | 6.6 | **0.001** |
| sex | 1 | 0.02 | 0.02 | 0.02 | 7.0 | 5.1 | **0.001** |
| environment | 1 | 0.01 | 0.01 | 0.01 | 3.6 | 3.3 | **0.001** |
| residuals | 110 | 0.24 | 0.00 | 0.32 | | | |
| total | 114 | 0.75 | | | | | |

*Note:*
Type II sum of squares is used. Significance (< 0.05) is indicated in bold.

Table S7: Effect of the environment (captive–wild difference) on lnCS evaluated by analysis of variance. Developmental phase, species, and sex are used as covariates.

| Source | $Df$ | $SS$ | $MS$ | $\eta^2$ | $F$ | $P$ |
|---|---|---|---|---|---|---|
| **Cranium** | | | | | | |
| Developmental phase | 1 | 1.983 | 1.983 | 0.722 | 959.7 | **< 0.001** |
| species | 1 | 0.226 | 0.226 | 0.109 | 109.2 | **< 0.001** |
| sex | 1 | 0.119 | 0.119 | 0.058 | 57.4 | **< 0.001** |
| environment | 1 | 0.023 | 0.023 | 0.010 | 11.3 | **0.001** |
| residuals | 119 | 0.246 | 0.002 | | | |
| **Mandible** | | | | | | |
| Developmental phase | 1 | 3.499 | 3.499 | 0.743 | 855.2 | **< 0.001** |
| species | 1 | 0.369 | 0.369 | 0.101 | 90.3 | **< 0.001** |
| sex | 1 | 0.145 | 0.145 | 0.041 | 35.4 | **< 0.001** |
| environment | 1 | 0.027 | 0.027 | 0.006 | 6.6 | **0.012** |
| residuals | 110 | 0.450 | 0.004 | | | |
| **Maxillary sinus** | | | | | | |
| Developmental phase | 1 | 7.871 | 7.871 | 0.545 | 248.8 | **< 0.001** |
| species | 1 | 1.585 | 1.585 | 0.209 | 50.1 | **< 0.001** |
| sex | 1 | 0.288 | 0.288 | 0.021 | 9.1 | **0.003** |
| environment | 1 | 0.156 | 0.156 | 0.009 | 4.9 | **0.028** |
| residuals | 121 | 3.827 | 0.032 | | | |

*Note:*
Type II sum of squares is used. Significance (< 0.05) is indicated in bold.

Table S8: Mean admixture proportion for each hybrid class in the Oike population.

| Hybrid class | Sex | $N$ | Autosome | X-chromosome | Y-chromosome | mtDNA |
|---|---|---|---|---|---|---|
| P0 | female | 13 | 0.00 | 0.00 | | 0.00 |
| | male | 10 | 0.00 | 0.07 | 0.00 | 0.00 |
| BC0 | female | 28 | 0.18 | 0.23 | | 0.00 |
| | male | 42 | 0.20 | 0.30 | 0.17 | 0.00 |
| F1 | female | 16 | 0.47 | 0.53 | | 0.00 |
| | male | 5 | 0.48 | 0.00 | 1.00 | 0.00 |
| F2 | female | 37 | 0.41 | 0.41 | | 0.00 |
| | male | 34 | 0.40 | 0.60 | 0.35 | 0.00 |
| BC1 | female | 31 | 0.77 | 0.85 | | 0.00 |
| | male | 33 | 0.76 | 0.48 | 1.00 | 0.00 |
| P1 | male | 11 | 1.00 | 1.00 | 1.00 | 1.00 |
| unknown | female | 7 | 0.66 | 0.55 | | 0.00 |
| | male | 14 | 0.61 | 0.49 | 0.54 | 0.00 |

*Note:*
The mean admixture proportions of autosomes and X chromosomes were calculated from the ADMIXTURE $Q$ score ($K$ = 2). See also Figure S2.

Table S9: Sex ratio for each hybrid class in the Oike population evaluated by chi-square test.

| Hybrid class | Female | Male | $\chi^2$ | $Df$ | $P$ | $P_{adj}$ † |
|---|---|---|---|---|---|---|
| P0 | 13 | 10 | 0.39 | 1 | 0.532 | 1.000 |
| BC0 | 28 | 42 | 2.80 | 1 | 0.094 | 0.660 |
| F1 | 16 | 5 | 5.76 | 1 | **0.016** | 0.115 |
| F2 | 37 | 34 | 0.13 | 1 | 0.722 | 1.000 |
| BC1 | 31 | 33 | 0.06 | 1 | 0.803 | 1.000 |
| P1 | 0 | 11 | 11.00 | 1 | **< 0.001** | **0.006** |
| unknown | 7 | 14 | 2.33 | 1 | 0.127 | 0.886 |

*Note:*
Significance (< 0.05) is indicated in bold.
† Bonferroni adjusted $P$ value.

Table S10: Age structure differences among hybrid classes in the Oike population evaluated by Fisher's exact test. The lower triangle shows $P$ values, while the upper Bonferroni-adjusted $P$ values.

| | P0 | BC0 | F1 | F2 | BC1 | unknown |
|---|---|---|---|---|---|---|
| P0 | | 1.000 | 1.000 | **< 0.001** | 1.000 | 0.601 |
| BC0 | 0.219 | | 0.143 | 1.000 | 1.000 | 1.000 |
| F1 | 0.356 | **0.010** | | **< 0.001** | 0.125 | 0.325 |
| F2 | **< 0.001** | 0.104 | **< 0.001** | | **0.030** | 1.000 |
| BC1 | 0.325 | 0.540 | **0.008** | **0.002** | | 1.000 |
| unknown | **0.040** | 0.749 | **0.022** | 0.520 | 0.431 | |

*Note:*
Significance (< 0.05) is indicated in bold.

Table S11: Morphological differences among hybrid classes at each developmental phase evaluated by analysis of variance. Sex is used as a covariate. $\eta^2$ indicates the effect size of the hybrid class. $P$-values are Bonferroni adjusted.

| | | Developmental phase | | | | | | | |
|---|---|---|---|---|---|---|---|---|---|
| | | 2 | 3 | 4 | 5 | 6 | 7 | 9 | 10 |
| **Cranium** | | | | | | | | | |
| lnCS | $\eta^2$ | 0.653 | 0.464 | 0.599 | 0.450 | 0.517 | 0.171 | 0.118 | 0.336 |
| | $P$ | < **0.001** | < **0.001** | 0.077 | 1.000 | 0.249 | 0.157 | 0.572 | < **0.001** |
| PC1 | $\eta^2$ | 0.213 | 0.099 | 0.223 | 0.155 | 0.368 | 0.042 | 0.104 | 0.173 |
| | $P$ | 0.070 | 1.000 | 1.000 | 1.000 | 1.000 | 1.000 | 1.000 | **0.006** |
| PC2 | $\eta^2$ | 0.677 | 0.376 | 0.718 | 0.695 | 0.355 | 0.545 | 0.673 | 0.603 |
| | $P$ | < **0.001** | **0.006** | **0.003** | 0.132 | 1.000 | **0.002** | **0.046** | < **0.001** |
| PC3 | $\eta^2$ | 0.164 | 0.493 | 0.324 | 0.531 | 0.620 | 0.310 | 0.426 | 0.482 |
| | $P$ | 0.403 | < **0.001** | 1.000 | 1.000 | **0.028** | 0.322 | 0.650 | < **0.001** |
| **Mandible** | | | | | | | | | |
| lnCS | $\eta^2$ | 0.564 | 0.410 | 0.540 | 0.559 | 0.464 | 0.282 | 0.154 | 0.335 |
| | $P$ | < **0.001** | **0.001** | 0.277 | 1.000 | 0.578 | **0.024** | 0.331 | < **0.001** |
| PC1 | $\eta^2$ | 0.267 | 0.066 | 0.426 | 0.481 | 0.215 | 0.013 | 0.177 | 0.157 |
| | $P$ | **0.013** | 1.000 | 0.723 | 1.000 | 1.000 | 1.000 | 1.000 | 0.069 |
| PC2 | $\eta^2$ | 0.403 | 0.710 | 0.758 | 0.642 | 0.673 | 0.615 | 0.711 | 0.639 |
| | $P$ | < **0.001** | < **0.001** | **0.005** | 0.486 | **0.026** | < **0.001** | **0.024** | < **0.001** |
| PC3 | $\eta^2$ | 0.127 | 0.242 | 0.166 | 0.044 | 0.431 | 0.061 | 0.299 | 0.128 |
| | $P$ | 0.709 | 0.148 | 1.000 | 1.000 | 0.516 | 1.000 | **0.023** | **0.002** |
| **Maxillary sinus** | | | | | | | | | |
| lnCS | $\eta^2$ | 0.315 | 0.365 | 0.617 | 0.611 | 0.559 | 0.655 | 0.696 | 0.762 |
| | $P$ | **0.002** | **0.008** | 0.060 | 0.343 | 0.085 | < **0.001** | **0.028** | < **0.001** |
| log ratio† | $\eta^2$ | 0.433 | 0.514 | 0.693 | 0.699 | 0.667 | 0.757 | 0.597 | 0.840 |
| | $P$ | < **0.001** | < **0.001** | **0.013** | 0.165 | **0.015** | < **0.001** | **0.025** | < **0.001** |

*Note:*
Significance (< 0.05) is indicated in bold.
† lnCS (sinus) - lnCS (cranium)

Table S12: Phenotypic trajectory analysis comparison results. Pairwise absolute differences in path distances (magnitude), pairwise correlations between trajectories (direction), and pairwise trajectory shape differences (shape) were evaluated.

| | | Magnitude | | Direction | | Shape | |
|---|---|---|---|---|---|---|---|
| Region | Comparison | *d* | *P* | *angle* | *P* | *d* | *P* |
| **Female** | | | | | | | |
| cranium | BC0:F2 | 0.006 | 0.862 | 14.50 | 0.224 | 0.21 | 0.137 |
| | BC0:BC1 | 0.000 | 0.983 | 11.20 | 0.441 | 0.20 | 0.143 |
| | BC0:fuscata | 0.026 | 0.317 | 13.27 | 0.080 | 0.08 | 0.572 |
| | F2:BC1 | 0.005 | 0.830 | 13.92 | 0.078 | 0.01 | 0.987 |
| | F2:fuscata | 0.020 | 0.413 | 14.16 | **0.029** | 0.13 | 0.285 |
| | BC1:fuscata | 0.025 | 0.185 | 10.14 | 0.067 | 0.12 | 0.283 |
| mandible | BC0:F2 | 0.013 | 0.685 | 12.30 | 0.812 | 0.09 | 0.586 |
| | BC0:BC1 | 0.004 | 0.886 | 9.83 | 0.907 | 0.12 | 0.449 |
| | BC0:fuscata | 0.028 | 0.283 | 12.67 | 0.394 | 0.02 | 0.970 |
| | F2:BC1 | 0.009 | 0.729 | 11.58 | 0.696 | 0.06 | 0.732 |
| | F2:fuscata | 0.015 | 0.541 | 17.36 | **0.038** | 0.07 | 0.656 |
| | BC1:fuscata | 0.024 | 0.221 | 11.82 | 0.096 | 0.10 | 0.440 |
| **Male** | | | | | | | |
| cranium | cyclopis_P0:BC0 | 0.017 | 0.558 | 9.40 | 0.308 | 0.06 | 0.708 |
| | cyclopis_P0:BC1 | 0.024 | 0.405 | 11.70 | 0.118 | 0.15 | 0.285 |
| | cyclopis_P0:fuscata | 0.014 | 0.578 | 13.88 | **0.001** | 0.17 | 0.185 |
| | BC0:BC1 | 0.041 | 0.151 | 10.14 | 0.186 | 0.18 | 0.082 |
| | BC0:fuscata | 0.003 | 0.888 | 10.40 | **0.015** | 0.19 | **0.028** |
| | BC1:fuscata | 0.038 | 0.126 | 12.09 | **0.016** | 0.03 | 0.815 |
| mandible | cyclopis_P0:BC0 | 0.030 | 0.264 | 10.24 | 0.520 | 0.14 | 0.368 |
| | cyclopis_P0:BC1 | 0.009 | 0.752 | 8.53 | 0.918 | 0.17 | 0.266 |
| | cyclopis_P0:fuscata | 0.023 | 0.342 | 9.71 | 0.472 | 0.16 | 0.259 |
| | BC0:BC1 | 0.039 | 0.147 | 9.55 | 0.677 | 0.31 | **0.009** |
| | BC0:fuscata | 0.007 | 0.759 | 12.31 | **0.018** | 0.30 | **0.005** |
| | BC1:fuscata | 0.033 | 0.186 | 10.58 | 0.302 | 0.01 | 0.980 |

*Note:*
Significance (< 0.05) is indicated in bold.

Table S13: Sidak-adjusted $P$-values of pairwise differences in slope (upper triangle) and elevation (lower triangle) of reduced major axis regression of maxillary sinus lnCS on cranial lnCS.

| | cyclopis_P0 | BC0 | F1 | F2 | BC1 | fuscata |
|---|---|---|---|---|---|---|
| **Female** | | | | | | |
| cyclopis_P0 | | 0.763 | 0.988 | 0.895 | 0.264 | **< 0.001** |
| BC0 | 1.000 | | 0.241 | 1.000 | 0.992 | **0.001** |
| F1 | 0.228 | 0.997 | | 0.310 | 0.089 | **0.003** |
| F2 | 0.089 | **< 0.001** | 0.999 | | 0.999 | **0.019** |
| BC1 | **< 0.001** | **< 0.001** | **< 0.001** | 0.083 | | 0.052 |
| fuscata | **< 0.001** | **< 0.001** | **< 0.001** | **< 0.001** | **< 0.001** | |
| **Male** | | | | | | |
| cyclopis_P0 | | 1.000 | 0.841 | 0.838 | 0.791 | 0.103 |
| BC0 | 0.972 | | 0.876 | 0.739 | 0.897 | 0.054 |
| F1 | **0.028** | 0.113 | | 0.323 | 1.000 | 0.214 |
| F2 | **0.003** | 0.089 | 1.000 | | 0.065 | 1.000 |
| BC1 | **< 0.001** | **< 0.001** | 1.000 | **0.034** | | **< 0.001** |
| fuscata | **< 0.001** | **< 0.001** | **< 0.001** | **< 0.001** | **< 0.001** | |

*Note:*
Significance (< 0.05) is indicated in bold.

### Supplementary Figures

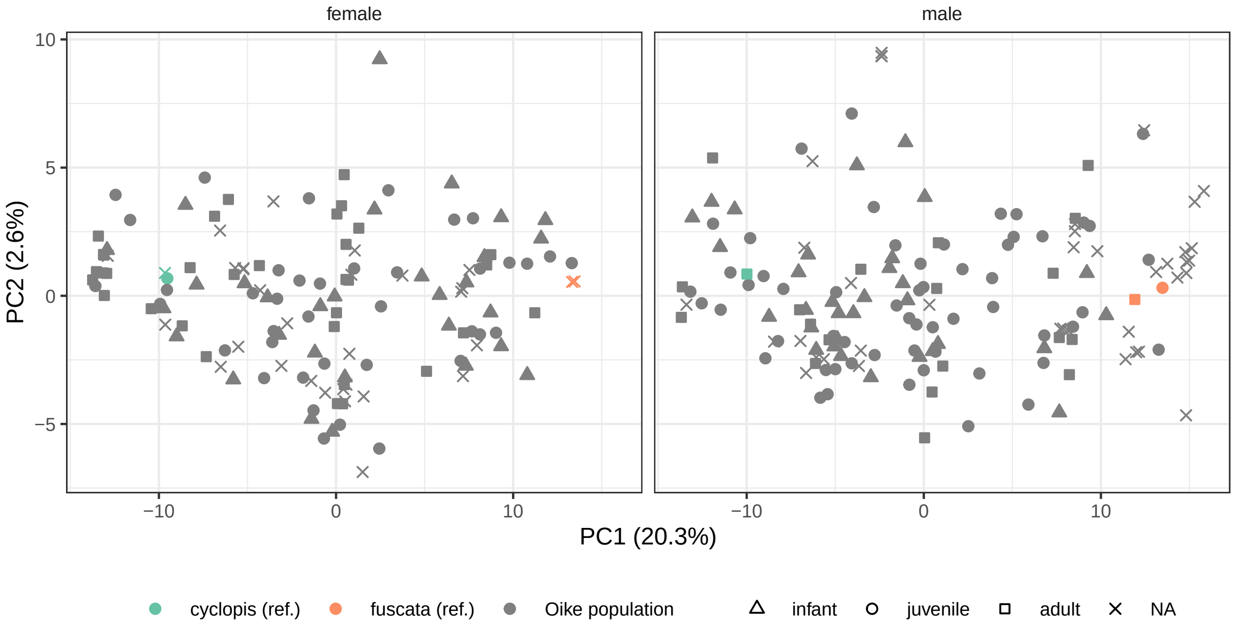


Figure S1: The principal component analysis of the autosomal SNPs. Blue indicates pure *Macaca cyclopis* housed at the Primate Research Institute, Kyoto University (Inuyama, Japan); magenta, pure *M. fuscata* housed at the Primate Research Institute; gray, individuals from the Oike population (Wakayama Prefecture, Japan). All samples were analyzed together, but each sex was plotted separately for better visualization.

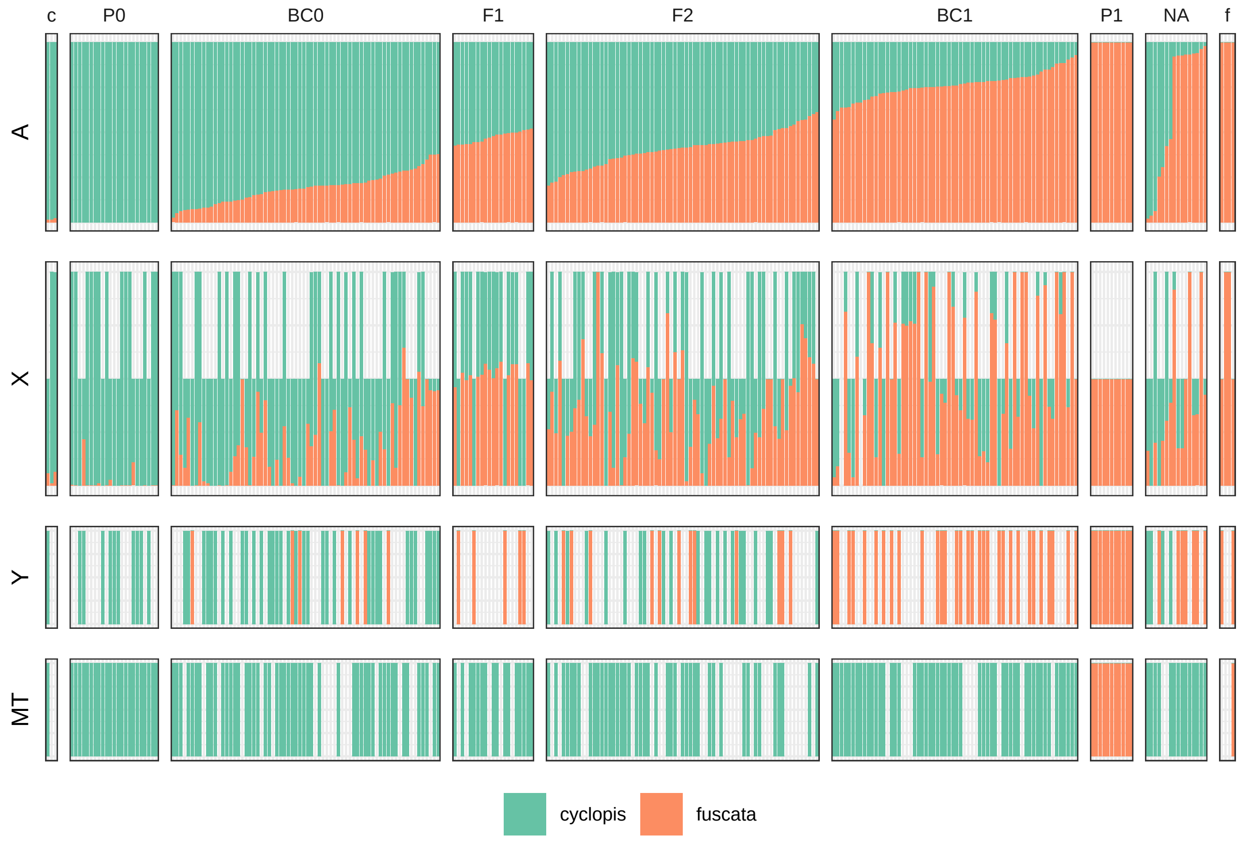


Figure S2: Ancestry panel blocked by hybrid class. The left (c) and right (f) blocks indicate the *Macaca cyclopis* and *M. fuscata* housed at the Primate Research Institute, Kyoto University (Inuyama, Japan), respectively; the middle blocks indicate the hybrid classes of the Oike population (Wakayama Prefecture, Japan). NA indicates an unknown hybrid class. Blue indicates *M. cyclopis*-type ancestry; magenta, *M. fuscata*-type ancestry; blank, missing data. Ancestry of autosomes (A) and X chromosomes (X) was estimated based on ddRAD SNPs using ADMIXTURE ($K = 2$). The ancestry of the male X chromosome was shown at half height. Y chromosome (Y) ancestry was estimated based on the $K$-means cluster ($K = 2$) of the first principal component score of ddRAD SNPs. Mitochondrial DNA (MT) ancestry was obtained from Kawamoto et al. (2008), supplemented by this study, where genotypes were identified by sequencing or restriction fragment length polymorphism.

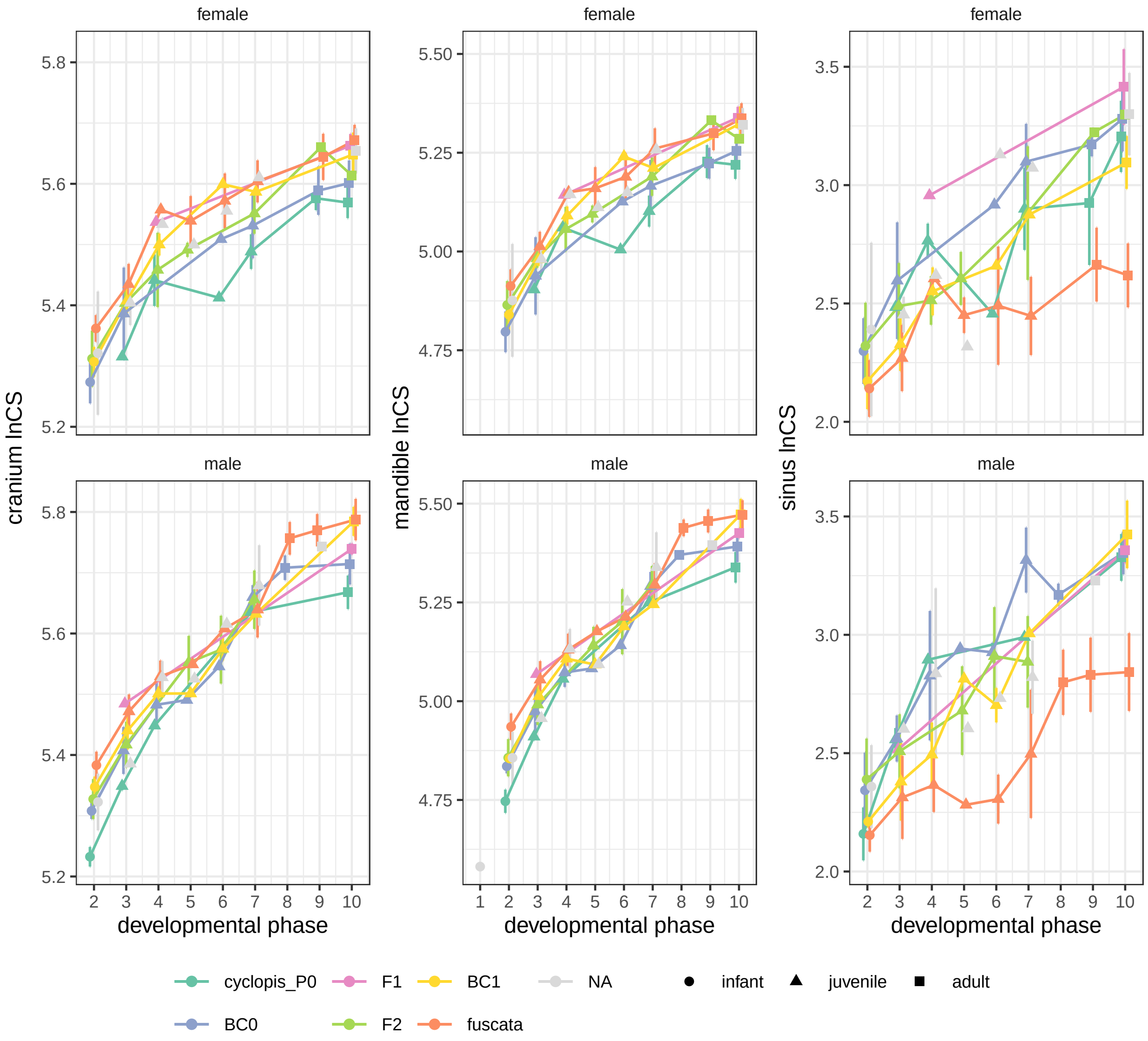


Figure S3: Developmental changes in the lnCS of the cranium, mandible, and maxillary sinus. Error bars are standard deviations.

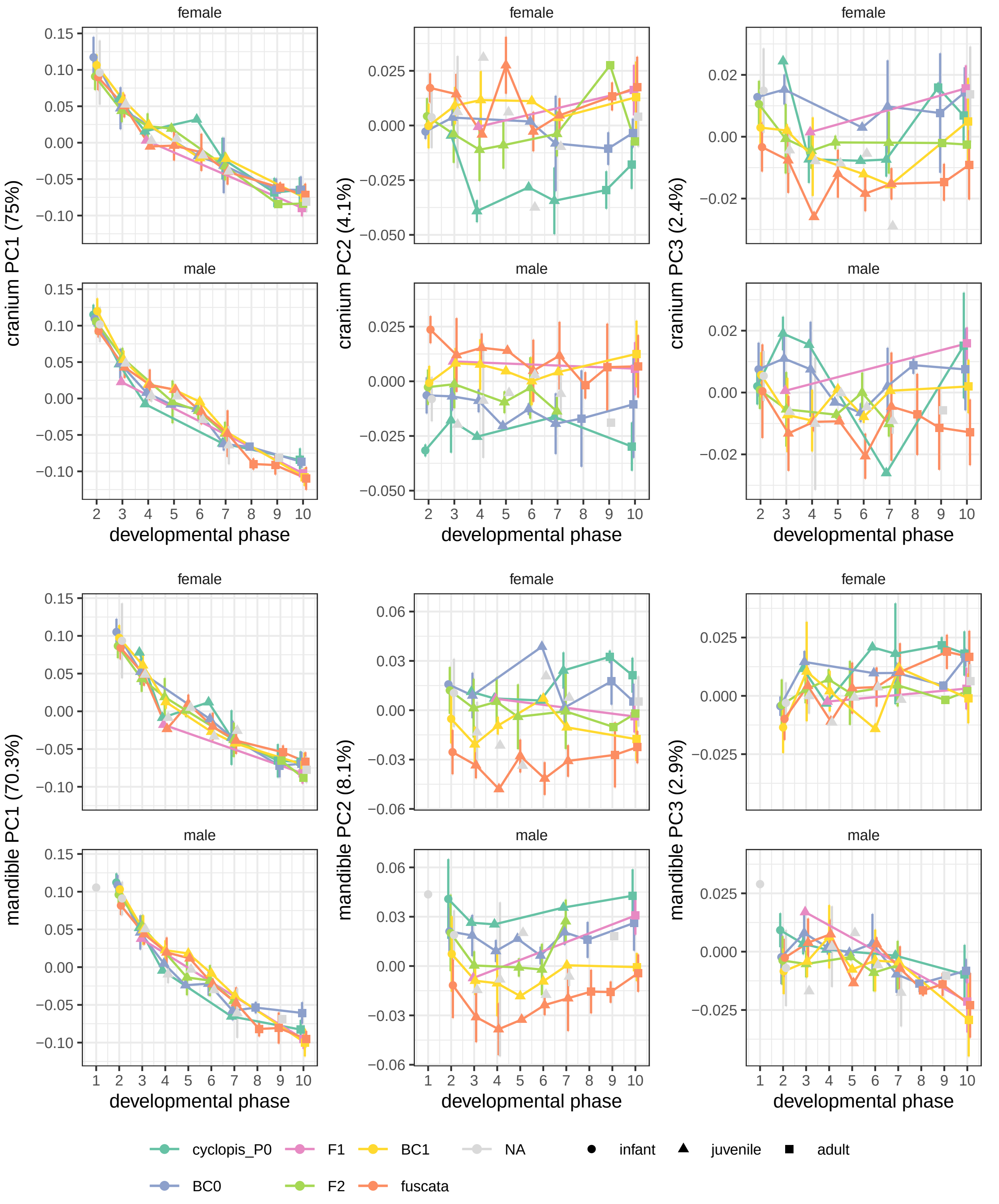


Figure S4: Developmental changes in cranial and mandibular shape. Error bars are standard deviations.

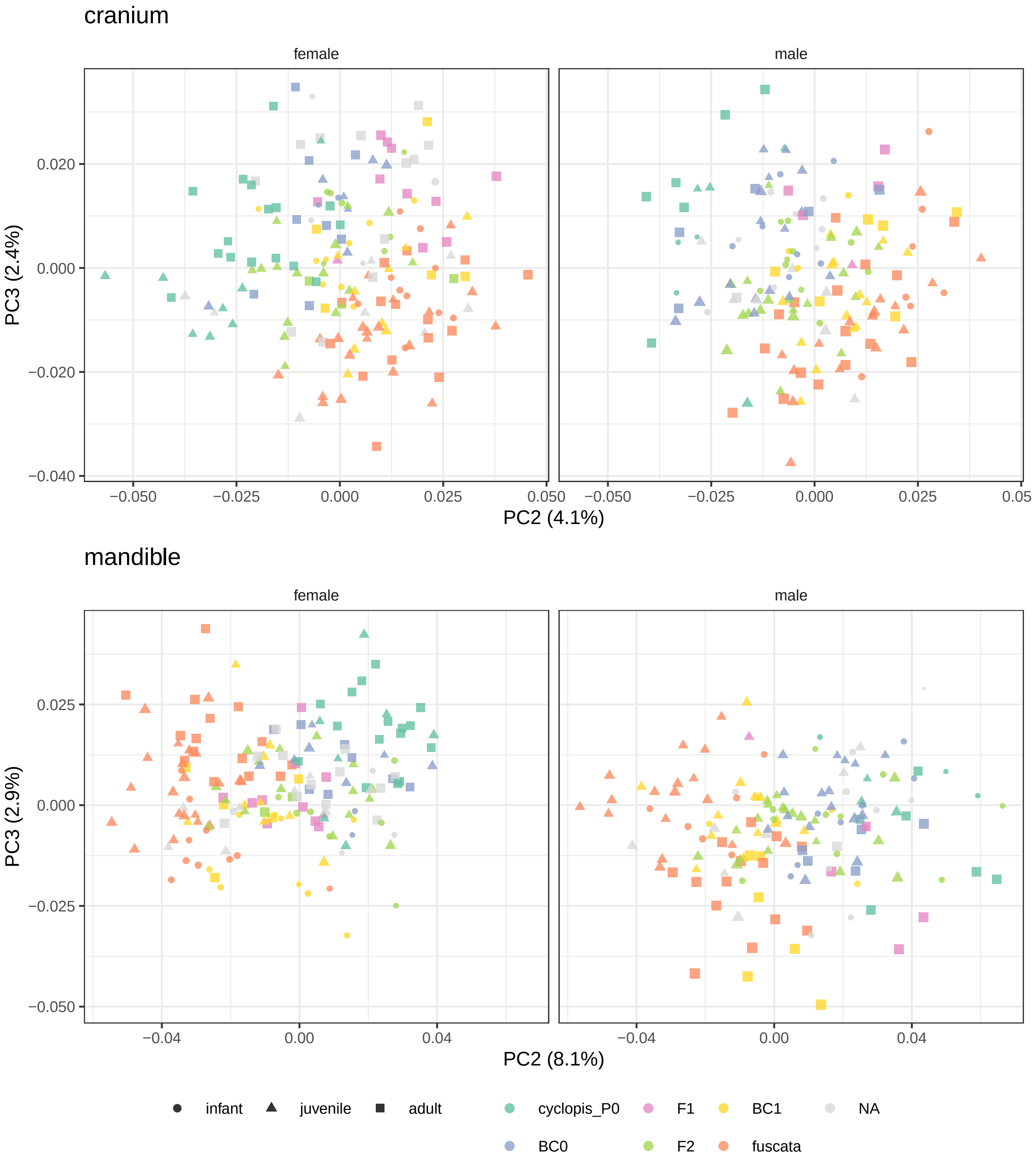


Figure S5: Variation in cranium and mandible shape represented by the second and third principal components (PCs). All samples were analyzed together, but each sex was plotted separately for better visualization. Points are scaled according to lnCS.

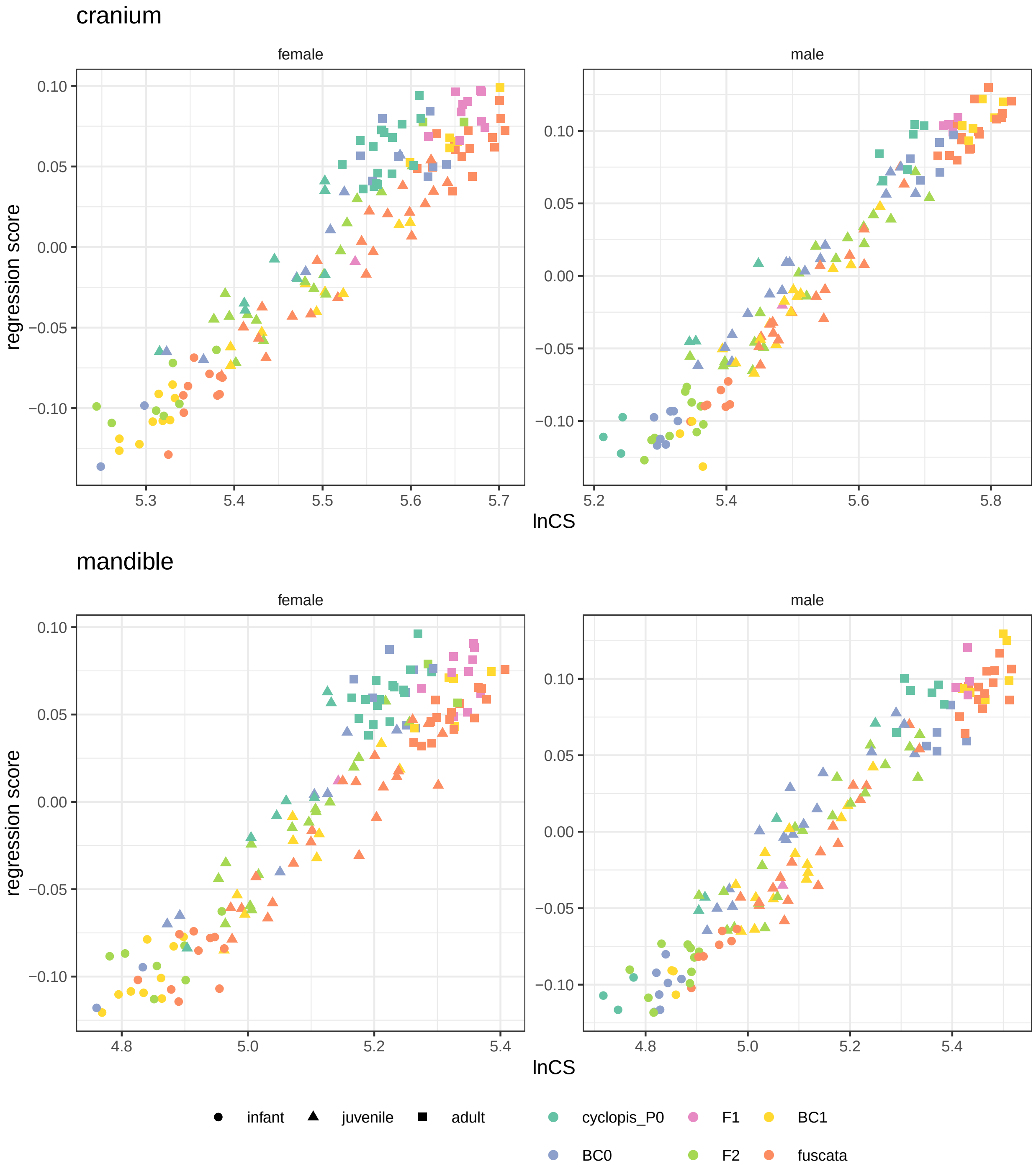


Figure S6: Ontogenetic allometry of cranial and mandibular shape represented by multivariate regression of shape against lnCS, hybrid class, and their interactions. Each sex was analyzed separately. A regression score is a projection of data on the normalized vector that expresses the covariation between shape and the regression coefficients for size, conditioned on other model effects (Adams et al. 2021).

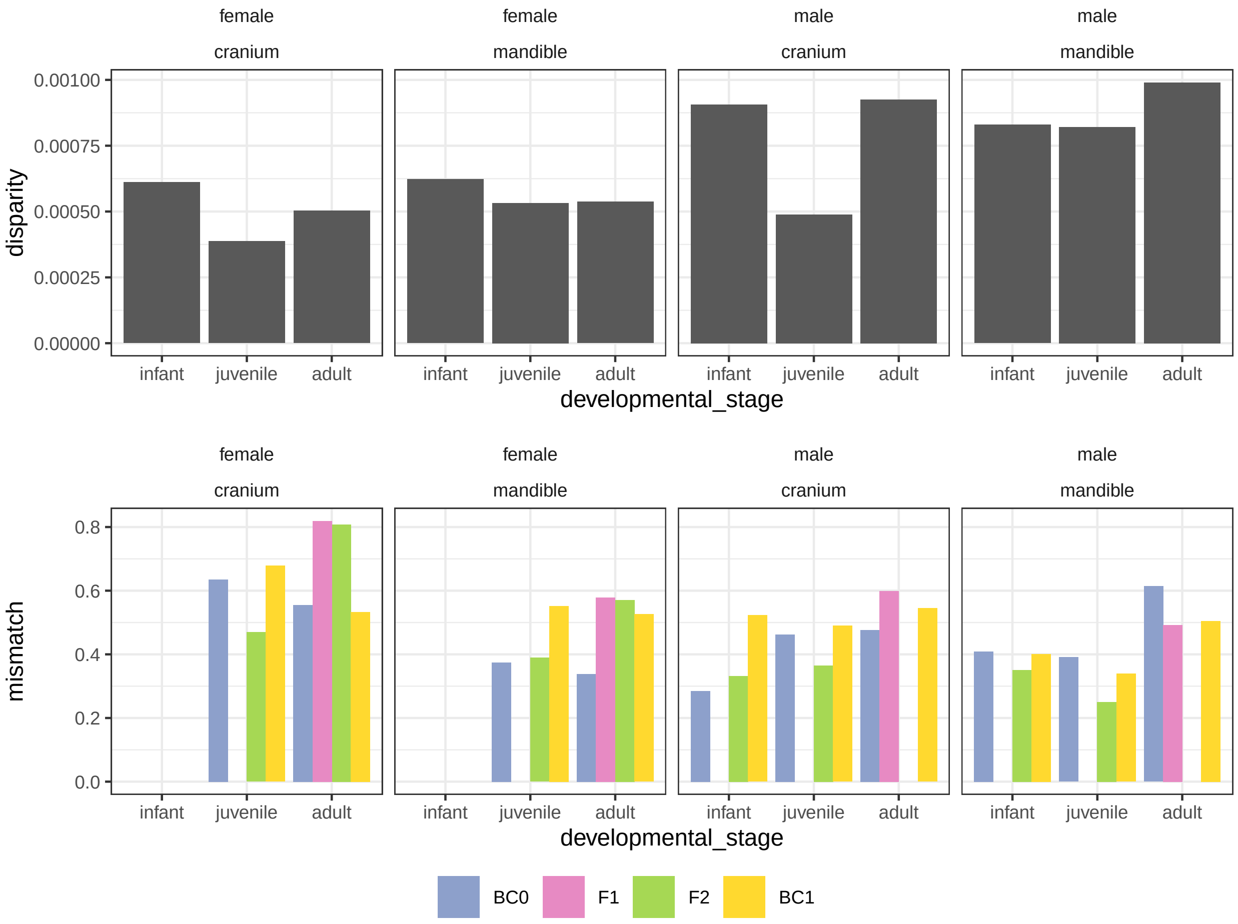


Figure S7: Developmental changes in disparity and mismatch (transgression). For disparity, hybrid classes with few samples in any of the three developmental stages (cyclopis_P0 and F1 for females; F1 and F2 for males) were omitted from the analyses. For mismatch, infant (all hybrid classes) and F1 juvenile data are missing for females, while F1 infant, F1 juvenile, and F2 adult data are missing for males.

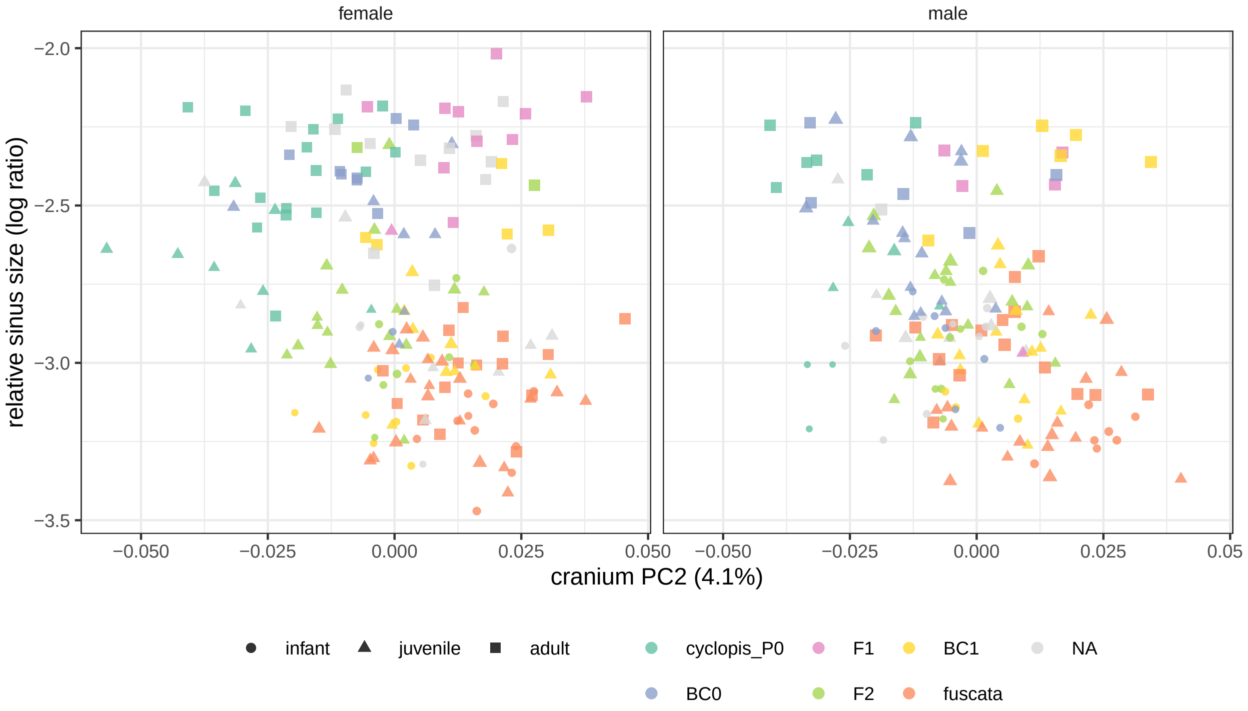


Figure S8: Cranium PC2 versus the relative size of the maxillary sinus (log ratio; sinus lnCS - cranium lnCS).
